# The RNA binding proteins LARP4A and LARP4B promote sarcoma and carcinoma growth and metastasis

**DOI:** 10.1101/2023.04.11.536377

**Authors:** Jennifer C. Coleman, Luke Tattersall, Val Yianni, Laura Knight, Hongqiang Yu, Sadie Hallett, Philip Johnson, Ana Caetano, Charlie Cosstick, Anne Ridley, Alison Gartland, Maria R Conte, Agamemnon E. Grigoriadis

## Abstract

RNA-binding proteins (RBPs) are emerging as important regulators of pathogenesis, including cancer. Here we reveal that the recently characterised RBPs LARP4A and LARP4B are differentially overexpressed in primary osteosarcoma and osteosarcoma lung metastases, as well as in prostate cancer. Depletion of LARP4A and LARP4B inhibited primary tumour growth and metastatic spread in xenograft studies, as well as inhibiting cell proliferation, motility and migration. Transcriptomic profiling combined with high content multiparametric cell cycle analysis unveiled a central role for LARP4B, but not LARP4A, in regulating cell cycle progression in osteosarcoma and prostate cancer cell lines, potentially through modulating the post-transcriptional regulation of RNA targets that include key cell cycle proteins such as Cyclins B1 and E2, Aurora B and E2F1. Our work assigns new functions to LARP4A and LARP4B as pro-tumorigenic proteins in bone and prostate cancer, highlights their similarities while indicating distinct functional aspects, and adds significantly to the rapidly increasing roles of RBPs in different cancer models. Uncovering clear biological roles for these paralogous proteins provides new avenues for identifying novel tissue-specific targets and potential druggable intervention.

## INTRODUCTION

Cancer is a heterogeneous disease in which a combination of genetic mutations and epigenetic changes enable cancer cells to divide more frequently than surrounding non-cancer cells, avoid cell death and spread to secondary sites as metastases.^1–2^ Changes in gene expression are central to the development, progression and spread of neoplastic disease. RNA binding proteins (RBPs) play central roles in this process, orchestrating the function and fate of coding and non-coding RNA molecules in a myriad of ways, including control of RNA splicing, polyadenylation, stability, localisation, transport, translation, processing, metabolism and decay.^3–6^ Growing evidence shows that RBPs are often dysregulated in cancer: their altered expression and/or mutations, as well as perturbations of RBP-RNA networks have been causally associated with cancer onset, progression and metastasis,^7–10^ but little is known of the mechanistic bases underlying these effects.

The La-Related Proteins (LARPs) are an ancient superfamily of RBPs with fundamental roles in RNA biology.^11^ They are present in all eukaryotes and consist of ∼250 members evolutionarily divided into 5 families (LARP1, LARP3, LARP4, LARP6 and LARP7). The hallmark of the LARPs is a conserved winged helix domain called the La motif (LaM) that, with a few exceptions, is associated with a downstream RNA-recognition motif (RRM1), together comprising the La-module.^11–13^ The La-module functions as a bipartite RNA recognition unit,^14–18^ albeit some exceptions have started to emerge.^19, 20^ Beyond this, each LARP family is characterised by family-specific domains and motifs implicated in RNA and/or protein partner interactions, using an assortment of binding modes and contributing to the diversification of functional profiles.^11, 13^ Although much remains to be explored, common as well as individual functional traits across the LARP families have started to be delineated.^11, 13^ Several LARPs have been associated with cancer,^10, 11, 21^ including the two human LARP4 paralogs, LARP4A and LARP4B, also referred to as LARP4 and LARP5, respectively. LARP4A and LARP4B have been described as predominantly cytoplasmic proteins that share overall domain organisation and display 74% sequence identity in their La-module and ∼40% throughout.^11^ They interact with the Poly(A) Binding protein (PABPC1) via a PAM2w motif located in the N-terminal region and a less-well defined PABPC-binding motif (PBM) downstream of the RRM1.^11, 20–25^ Their common features also include interaction with receptor for activated kinase C (RACK1), enhancement of translation and mRNA stability, association with ribosomes and localisation to stress granules.^11, 23–25^

Beyond this, other functions of either LARP4A or LARP4B have been steadily emerging. For instance, LARP4A has been shown to exhibit protection of mRNA 3’ poly(A) tail lengths and the associated mRNA stabilisation,^26, 27^ contribute to mRNA metabolism during activation of T cells immune response,^28^ play roles in the gene regulatory network governed by metazoan miRNAs^29^ and affect local mRNA translation at mitochondria level,^30^ while LARP4B appears to reduce cell and organ sizes in *Drosophila melanogaster* via dMyc regulation.^31^ These findings may possibly underline functional differences between LARP4A and LARP4B, thereby endorsing a sub- or neo-functionalisation hypothesis for these two paralogs that have been maintained through evolution.^32, 33^ However, notably, data remain patchy and limited, as investigations have not been pursued in parallel on both proteins, precluding a comprehensive comparative profiling of LARP4A and LARP4B’s functional capacities. To date, investigations of RNA targets and RNA binding preferences have revealed that LARP4A associates with poly(A) RNA primarily via the PAM2w motif in a manner that is mutually exclusive to PABPC MLLE interaction,^20, 24^ whereas LARP4B binds AU-rich sequences in 3’ untranslated regions (UTRs) in mature mRNAs^25^ using a different molecular mechanism (M.R.C., unpublished). How these different RNA recognition capabilities translate to the function of LARP4A and LARP4B remains to be elucidated.

The roles of LARP4A and LARP4B in cancer biology are equally not well understood, but sporadic evidence is accumulating in line with the increasing interest in RNA binding proteins in cancer progression.^10^ We and others first described LARP4A as a regulator of cancer cell morphology in a RNAi *Drosophila* screen and subsequently shown to affect cell migration in breast, prostate and ovarian cancer cell lines, albeit with varying effects.^34–36^ Since then, LARP4A is often coined a tumour suppressor, however its roles in tumorigenesis and cancer growth have not yet been elucidated. Work on LARP4B has revealed varying roles depending on tumour or tissue type: in glioma, LARP4B has tumour suppressive properties because of its growth-restrictive roles,^37, 38^ whereas it possesses oncogenic growth-promoting effects in an Acute Myeloid Leukaemia (AML) mouse model.^39^ Data from the TCGA TARGET GTEx study have associated altered *LARP4A* and *LARP4B* mRNA expression with different cancer subsets and, intriguingly, upregulation of *LARP4B* mRNA was suggested to be an independent risk factor for the prognosis of liver cancer patients.^40^ LARP4A and/or LARP4B have also appeared in several transposon mutagenesis screens to identify new genes driving tumorigenesis for colorectal cancer,^41^ hepatocellular carcinoma,^42^ malignant peripheral nerve sheath tumours^43^ and metastatic medulloblastoma.^44^ Furthermore, enrichment of nonsense mediated decay (NMD)-elicit mutations in LARP4B was revealed in hypermutated stomach adenocarcinoma.^45^

The increasing association between the LARP4 proteins and cancer suggests complex and context-dependent functional roles for LARP4A and LARP4B in tumour progression. In this study, we have determined the contributions of LARP4A and LARP4B to cancer progression using osteosarcoma and prostate cancer cells, representing two distinct cancer origins, mesenchymal and epithelial, respectively. Using loss- and gain-of-function approaches, we provide a systematic comparison of their similarities and differences with respect to their cellular, transcriptional and biological effects *in vivo* and *in vitro*. We reveal key roles of LARP4A and LARP4B in cancer progression and identify candidate mechanisms underlying these functions.

## RESULTS

### LARP4A and LARP4B are overexpressed in cancer tissues

To understand the possible functions of LARP4A and LARP4B as oncoproteins or tumour suppressors *in vivo*, we employed in the first instance an established osteosarcoma transgenic mouse model.^46, 47^ Immunohistochemical staining of primary tumours revealed high expression of both LARP4A and LARP4B proteins in transformed osteoblasts lining neoplastic bone as well as within the intratumoral stroma, with negligible levels in normal bone (Figures 1A and 1B). Further analysis of osteosarcoma lung metastasis^47^ detected high LARP4A expression within the normal pulmonary interstitium as well as in metastatic nodules (Figures 1C(i) and 1D(i,ii)), whereas LARP4B displayed a striking profile with very low levels in normal lung interstitial cells and a relatively higher expression in metastatic foci (Figures 1C(ii) and 1D(iv,v)). Interestingly, nuclear staining for both LARP4A and LARP4B was evident in subpopulations of transformed osteoblasts and tumour stromal cells in primary osteosarcoma as well as in lung metastases (Figures 1B(iii,vi) and 1D(iii,vi)).

**Figure 1.**
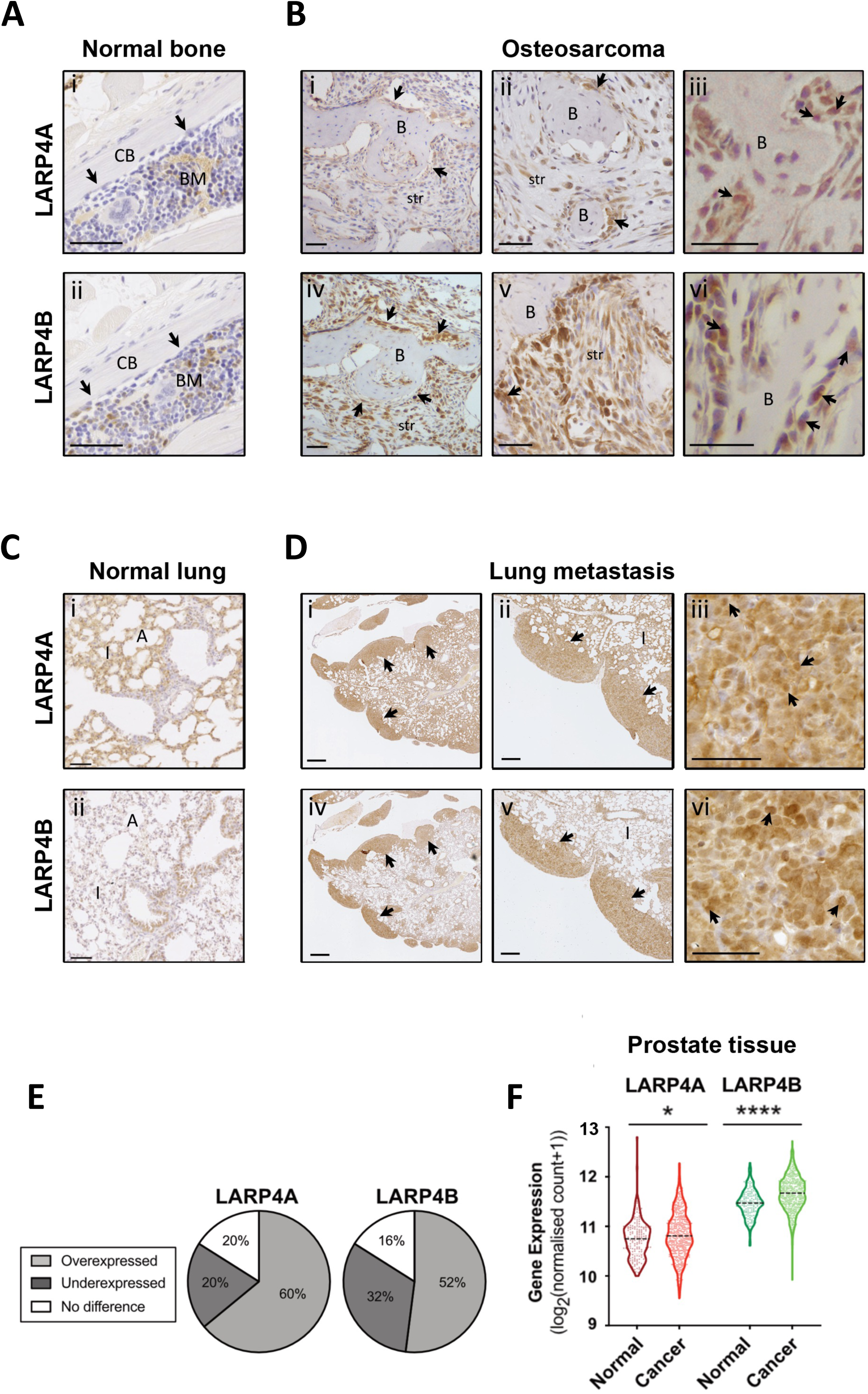
Expression of LARP4A and LARP4B in normal and cancer tissues. (A) Immunohistochemical analysis of LARP4A and LARP4B protein expression in normal murine bone. Control bones show little to no expression of either LARP4A (A(i)) or LARP4B (A(ii)) in osteoblasts (arrows). CB - cortical bone (normal bone); BM – bone marrow. Scale bars, 50µm. (B) Immunohistochemical analysis of LARP4A and LARP4B protein expression in a primary murine osteosarcoma^47, 48^. Increasing magnification of osteosarcoma tissues stained with LARP4A (B(i-iii)) or LARP4B (B(iv-vi)) show higher overall expression of LARP4B in transformed osteoblasts (arrows) and tumour stromal cells (str) compared to LARP4A. Arrows in B(iii) and B(vi) indicate nuclear staining of LARP4A and LARP4B, respectively. B – neoplastic bone (tumour). Scale bars, 50µm. (C) Immunohistochemical analysis of LARP4A and LARP4B protein expression in normal murine lung tissue. Control lungs show markedly higher expression of LARP4A (C(i)) in pneumocytes of the lung interstitium compared to LARP4B (C(ii)). Scale bars, 200µm. (D) Immunohistochemical analysis of LARP4A and LARP4B protein expression in lungs harbouring osteosarcoma metastatic nodules following tail vein injection. Increasing magnification of lungs stained with LARP4A (D(i-iii)) or LARP4B (D(iv-vi)) show high expression in metastatic nodules (arrows). Arrows in D(iii) and D(vi) indicate nuclear staining of LARP4A and LARP4B, respectively. I – lung interstitium; A – alveolar sacs. Scale bars, (D(ii,v)) 200µm; (D(i,iii,iv,vi) 500µm. (E) Percentage of tumour types where *LARP4A* or *LARP4B* are significantly overexpressed or underexpressed compared to control tissue according to TCGA TARGET GTEx mRNA expression analysis across 23 tissue types. Full statistical analysis across tumour types can be seen in Figure S1. (F) Comparison of *LARP4A* and *LARP4B* mRNA expression in normal prostate and prostate adenocarcinoma samples. Violin plots represent the mean expression in normal (n=152) or cancer (n=496) tissues according to the TGCA TARGET GTEx database. * p<0.05, ****p<0.0001. See also Figure S1.

Immunohistochemical staining of prostate cancer tissue also showed high expression of both LARP4A and LARP4B proteins (data not shown), indicating that these RNA binding proteins are likely to be highly expressed in cancer. This was endorsed following a pan-cancer analysis of LARP4A and LARP4B expression in patient tumour samples and normal tissues using the TCGA TARGET GTEx database^48^ which showed that *LARP4A* and *LARP4B* mRNA are significantly overexpressed in the majority (60% and 52% respectively) of the 25 human cancer subsets, and significantly underexpressed in 20% and 32% of these compared to normal tissue controls (Figures 1E, S1A, and S1B). Interestingly, both proteins were significantly upregulated in prostate adenocarcinoma tissue although a lack of normal sample availability likely prevented accurate quantification of expression in bone/sarcoma tissues (Figures 1F, S1A, and S1B).

### Depletion of LARP4A and LARP4B markedly inhibits tumour growth *in vivo*

To investigate the functional consequences of LARP4A/LARP4B in cancer progression, we performed xenograft studies following genetic silencing of each protein in representative osteosarcoma (MNNG/HOS; MG63) and prostate cancer (PC3) cell lines. Western blot and qPCR analyses indicated efficient silencing of LARP4A and LARP4B using targeted siRNAs in PC3, MNNG/HOS and MG63 cells (Figure 2A and data not shown). Subcutaneous injection of LARP4A/LARP4B-depleted PC3 and MNNG/HOS cells in immunocompromised mice showed a marked reduction in tumour volume by 50-85% in both PC3- and MNNG/HOS-derived xenografts (Figures 2B and 2C). Tumours formed by LARP4A and LARP4B-depleted cells had significantly lower expression of the mitosis and proliferation markers pHH3 and Ki67 while expression of the apoptotic marker Cleaved Caspase-3 was not altered (Figures 2D, 2E, and S2A-D), suggesting that LARP4A and LARP4B support prostate cancer and osteosarcoma tumour proliferation *in vivo*. We further assessed LARP4A and LARP4B tumorigenic capacities using a more biologically relevant orthotopic osteosarcoma xenograft model.^49^ Paratibial injection of MNNG/HOS cells depleted for either LARP4A or LARP4B resulted in a striking reduction in tumour formation compared to controls, as demonstrated by the low/undetectable luciferase luminescence signal over the experimental time-course and a significant decrease in tumour volume (Figures 2F and 2G). As MNNG/HOS cells are known to produce ectopic bone in tumour xenografts,^49^ we also monitored the bone-forming potential of LARP4A and LARP4B knockdown xenografts. Whilst the presence of ectopic bone was confirmed by microCT analysis in siControl xenografts, this was completely absent or dramatically reduced in bones engrafted with siLARP4A- and siLARP4B-depleted cells, respectively (Figure 2H). Histological analysis demonstrated large tumour masses formed by siControl cells including ectopic bone, both of which were absent or hugely diminished in siLARP4A and siLARP4B xenografts (Figure 2I).

**Figure 2.**
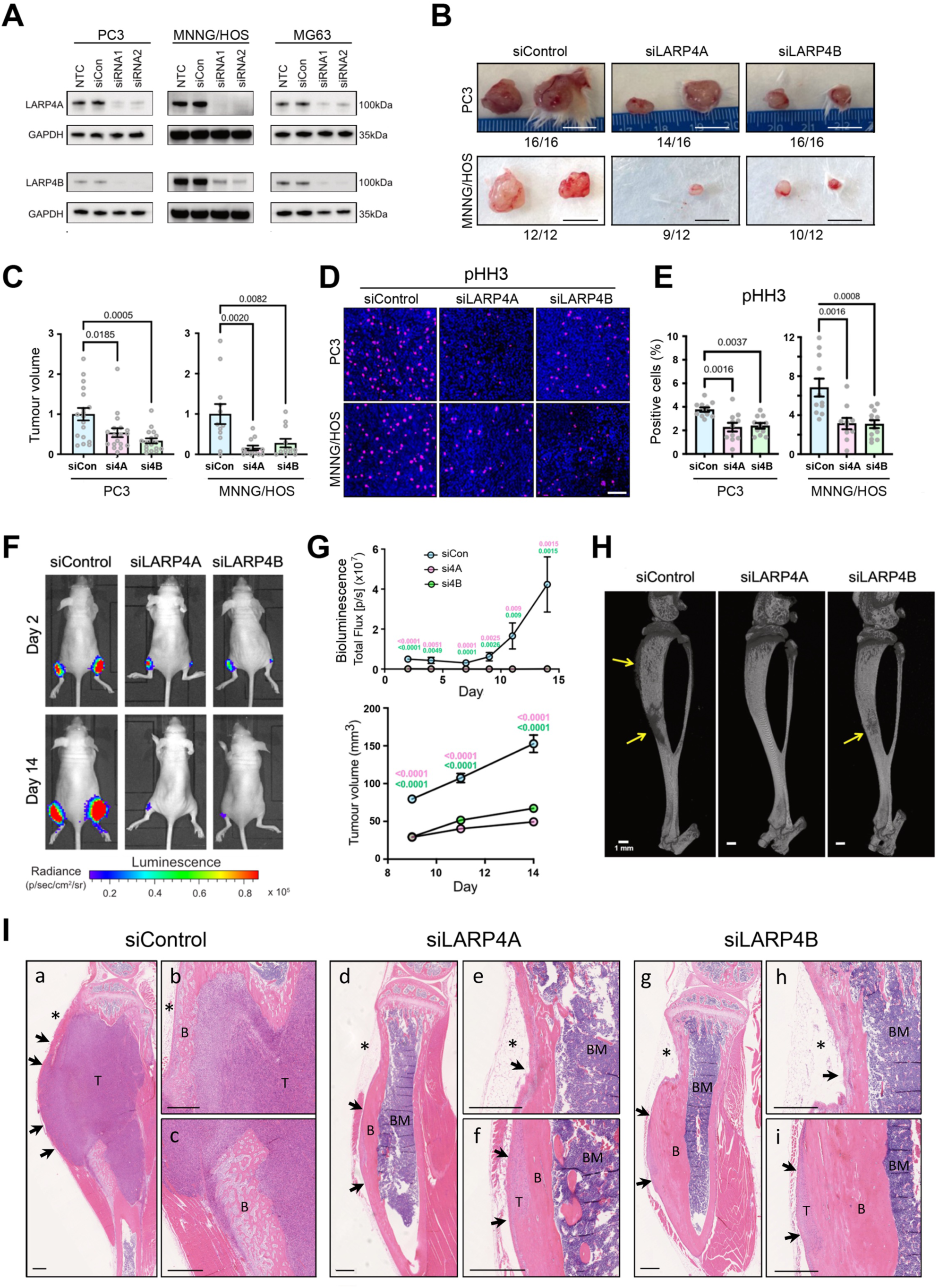
Depletion of LARP4A and LARP4B inhibits tumour formation *in vivo*. (A) Western blot analysis of endogenous LARP4A and LARP4B expression in PC3, MNNG/HOS and MG63 cells 72h following transfection with specific siRNAs targeting LARP4A (siLARP4Asi1/si2) or LARP4B (siLARP4Bsi1/si2) as indicated. Non-transfected (NTC) and non-targeting siRNA (siCon) controls are shown and GAPDH was used as a loading control. (B) Representative images of xenograft tumours formed following subcutaneous injection of siControl, siLARP4A or siLARP4B knockdown PC3 (top) and MNNG/HOS (bottom) cells. Numbers below each panel denote the proportion of engraftment of each cell line after 2wks. Scale bars, 1cm. (C) Quantification of volumes of subcutaneous tumours shown in (B) formed by siLARP4A and siLARP4B knockdown PC3 (*left*) and MNNG/HOS (*right*) cells, 2wks post-engraftment and normalised to siControl (siCon) tumours. The data represent the mean ± SEM, n=6-8 mice per condition. p-values are indicated. (D) Representative immunofluorescence images of pHH3 and DAPI staining in subcutaneous tumours formed by engraftment of PC3 or MNNG/HOS cells transfected with non-targeting siRNA controls (siControl) or siRNAs targeting LARP4A (siLARP4A) or LARP4B (siLARP4B). Scale bar, 100µm. (E) Quantification of the proportion of cells within PC3 (*left*) and MNNG/HOS (*right*) xenografts expressing pHH3 shown in (D), as a percentage the total number of cells (DAPI). Bars represent the mean ± SEM per tumour, n=9-16 individual tumours from 6-8 mice. p-values are indicated. (F) Representative bioluminescence images of paratibial xenografts of control (siControl) and LARP4A (siLARP4A) or LARP4B (siLARP4B) knockdown MNNG/HOS cells at 2d and 14d post-engraftment. (G) Quantification of the bioluminescence signal (*top*) and tumour volume (*bottom*) in mouse tibiae over a 14d time course. Data represent the mean ± SEM, two tibiae measured from n=8 mice per control or knockdown condition. p-values are indicated. (H) Representative microCT scans of murine tibiae following engraftment with control (siControl), LARP4A (siLARP4A) or LARP4B (siLARP4B) knockdown MNNG/HOS cells. Arrows highlight areas of malignant bone deposits as a result of osteosarcoma tumour formation. Scale bars, 1mm. (I) Representative histological images of tibiae 14d post-engraftment with siControl, siLARP4A or siLARP4B knockdown MNNG/HOS cells. Low magnification images show a clear reduction in tumour mass in both siLARP4A (I(d)) and siLARP4B (I(g)) knockdown xenografts compared to siControls (I(a)). High magnification images in panels I(b) and I(c) (siControl), I(e) and I(f) (siLARP4A) and I(h) and I(i) (siLARP4B) show areas of tumour tissue (T) that are markedly reduced in the LARP4A and 4B knockout xenografts. B, Bone, T, Tumour, BM, Bone Marrow. Asterisks indicate the cell injection sites. Arrows indicate the tumour perimeter which is reduced in the knockout xenografts. Scale bars, 500µm. See also Figure S2.

Finally, we assessed the metastatic potential of LARP4A and LARP4B-depleted osteosarcoma cells *in vivo*. Whereas siControl MNNG/HOS cells formed detectable nodules in lungs within 7d, LARP4A and LARP4B-depleted cells formed significantly less foci (Figures 3A-C, and S3), with a concurrent, significant decrease in their ability to seed or expand in pulmonary tissue (Figures 3D and 3E). This was confirmed with the additional osteosarcoma cell line MG63, which demonstrated a significant decrease in cell engraftment as early as 24h post-injection (Figures 3F and 3G), indicating an essential role for these proteins in promoting metastatic colonisation by tumour cells. Taken together, these *in vivo* expression and xenograft studies suggest clear pro-tumorigenic and metastatic roles for both LARP4A and LARP4B.

**Figure 3.**
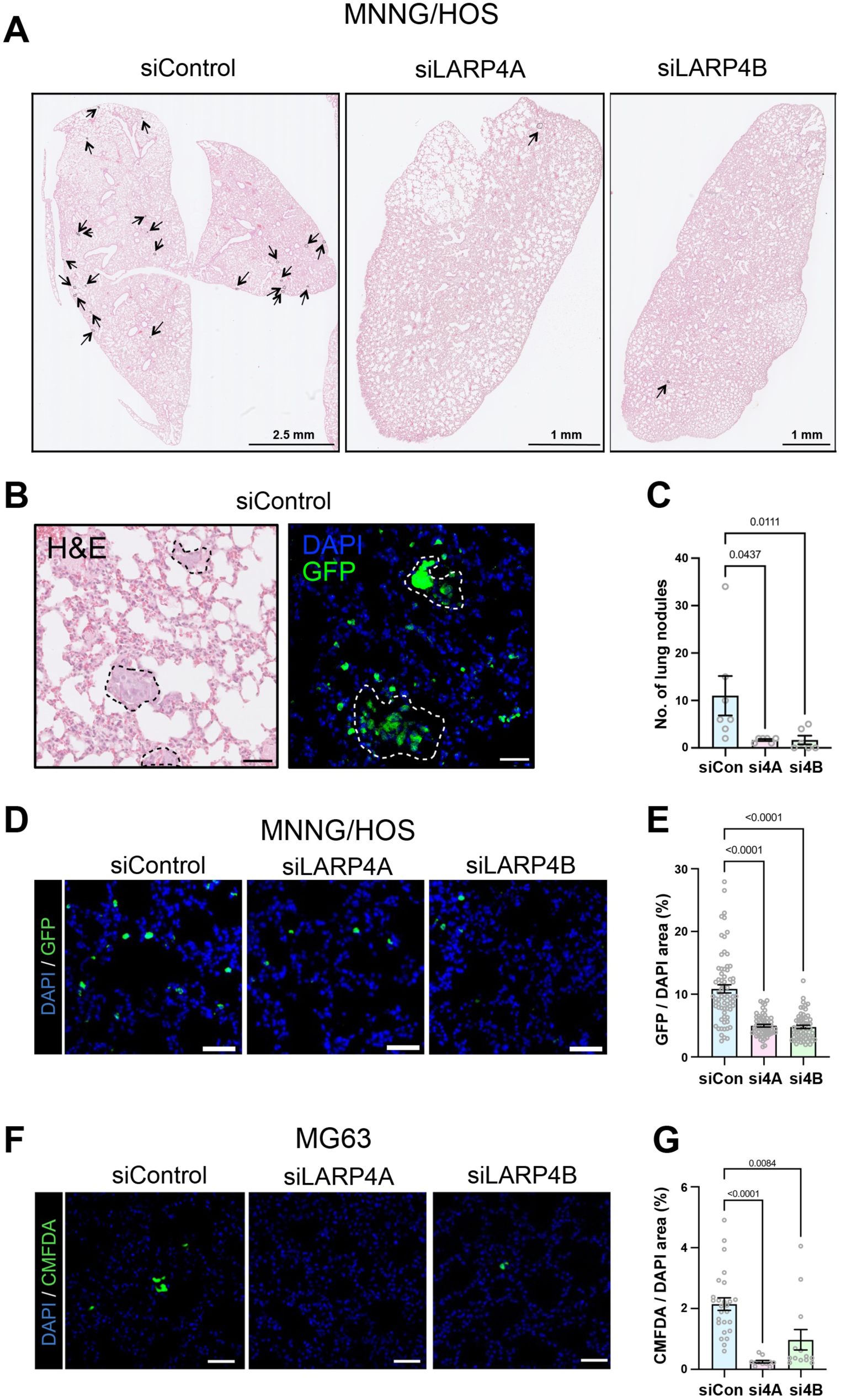
Depletion of LARP4A and LARP4B inhibits lung metastasis. (A) Low magnification histological images (H&E) of lungs 7d following tail vein injection of control (siControl), LARP4A (siLARP4A) or LARP4B (siLARP4B) knockdown MNNG/HOS cells. Black arrows indicate metastatic nodules. Scale bars denoted within images. (B) High magnification of representative histological (H&E) and immunofluorescence (GFP/DAPI) images of MNNG/HOS GFP-positive metastatic nodules 7d following tail vein injection. Scale bar, 50µm. (C) Quantification of the total number of metastatic nodules lesions per mouse 7d following tail vein injection of control (siCon), LARP4A (si4A) or LARP4B (si4B) knockdown MNNG/HOS cells. The data represent the mean ± SEM, n=6 mice per condition. p-values are indicated. (D) Representative immunofluorescence images of MNNG/HOS-GFP+ metastatic lung nodules 7d following tail vein injection of siControl, siLARP4A or siLARP4B knockdown cells. Scale bars, 50µm. (E) Quantification of the proportion of MNNG/HOS-GFP+ cells 7d following tail vein injection of knockdown cells shown in (D). The data represent the mean ± SEM, 10 randomly selected fields in n=6 lungs per condition. p-values are indicated. (F) Representative immunofluorescence images of murine lungs 24h following tail vein injection of CMFDA-labelled siControl, siLARP4A or siLARP4B knockdown MG63 cells. Scale bars, 50µm. (G) Quantification of the proportion of CMFDA-labelled MG63 cells 24h following tail vein injection of knockdown cells shown in (F). The data represent the mean ± SEM, n=3 mice per condition. p-values are indicated. See also Figure S3.

### LARP4A and LARP4B promote cell proliferation and LARP4B influences cell cycle progression

To uncover potential mechanisms underpinning the pro-tumorigenic roles of LARP4A and LARP4B *in vivo*, we compared the function of these paralogs on cell proliferation and viability. MTS assays demonstrated that depletion of either LARP4A or LARP4B proteins by targeted siRNA silencing significantly reduced the viability of PC3, MG63 and MNNG/HOS cell lines compared to controls, and this was verified by quantification of cell numbers (Figures 4A-D), suggesting that both proteins positively regulate cell proliferation.

**Figure 4.**
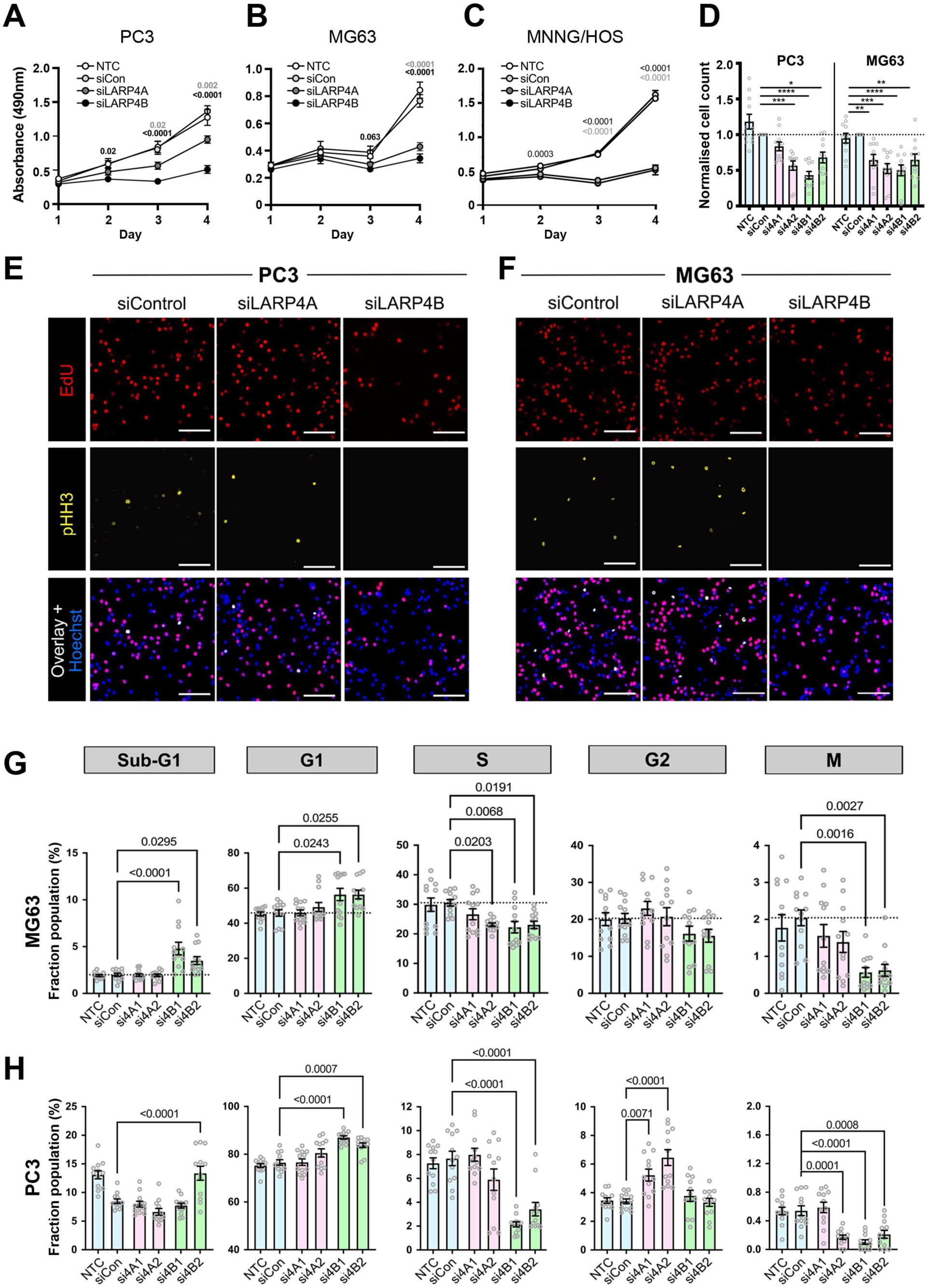
LARP4A and LARP4B regulate cell proliferation and LARP4B affects cell cycle progression. (A-C) MTS viability assays of PC3 (A), MG63 (B) and MNNG/HOS (C) control cells (non-transfected NTC, siCon) and siLARP4A or siLARP4B knockdown cells over a 4d time course. The data represent the mean ± SEM, n=8 from 4 independent experiments. p-values are indicated. (D) Quantification of PC3 (*left*) and MG63 (*right*) cell numbers after 72h following knockdown of siLARP4A (si4A1, si4A2) and siLARP4B (si4B1, si4B2) normalised to non-targeting siRNA (siCon) controls. NTC – non-transfected controls. The data represent the mean ± SEM, n=11 from three independent experiments. *p<0.05, **p<0.01, ***p<0.001, ****p<0.0001. (E-F) Immunofluorescence images of siControl, siLARP4A and siLARP4B knockdown PC3 (E) and MG63 (F) cells following EdU incorporation and pHH3 staining. Hoechst was used as counterstain. Scale bars, 200µm. (G-H) Multiparametric cell cycle analysis of PC3 (**G**) and MG63 (**H**) cells either non-transfected (NTC) or transfected with non-targeting control (siCon) and siRNAs targeting LARP4A (si4A1, si4A2), LARP4B (si4B1, si4B2) as indicated. The fraction of the population at each cell cycle stage (sub-G1, G1, S, G2 and M phases) for each cell type is depicted. The data represent the mean ± SEM, n=12 from four independent experiments. p-values are indicated. See also Figure S4.

Based on reports that LARP4 proteins, in particular LARP4B, might perturb cell cycle,^38, 39^ we investigated cell cycle distribution in LARP4A- and LARP4B-depleted MG63 and PC3 cells. High content multiparametric analysis^50^ of separate cell cycle stages showed that LARP4B knockdown in both cell lines significantly increased the fraction of cells in sub-G1 and G1 phases with a concomitant reduction in S and M phases (Figures 4E-H), indicating that depletion of LARP4B resulted in reduced cell cycle entry, slower progression through G1/G2 phases and decreased mitosis. In contrast, LARP4A depletion did not appreciably alter cell cycle (Figures 4E-H). These results were confirmed by overexpressing GFP-tagged proteins (Figures S4A and S4B). LARP4B overexpression increased the fraction of cells in S and M phases in both MG63 and PC3, whereas LARP4A overexpression had no effect (Figures S4C and S4D). LARP4B therefore enhances cell cycle progression, possibly by promoting cellular replication and mitosis. In agreement with this, RNAseq analysis revealed that siRNA-depletion of LARP4B in MG63 and PC3 cells resulted in decreased RNA levels for genes over-represented in biological processes involved in regulation of mitosis, spindle organisation and cell cycle phase transition (Figures 5A and 5B). Specific genes include cyclins (*CCNB1 and CCNE2),* the positive cell-cycle regulator *CDC7,*^51^ the transcription factor *E2F1* which promotes the expression of genes involved in S-phase entry and mitosis,^52^ the proliferation marker *KI67,* whose expression is controlled during cell cycle^53^ and *BRCA2,* the tightly regulated expression of which is highly critical in proliferation and cell cycle checkpoints (Figure 5B).^54^ Intriguingly, and in agreement with cell cycle multiparametric analysis, these changes were not observed in LARP4A-depleted cells (Figure 5B).

**Figure 5.**
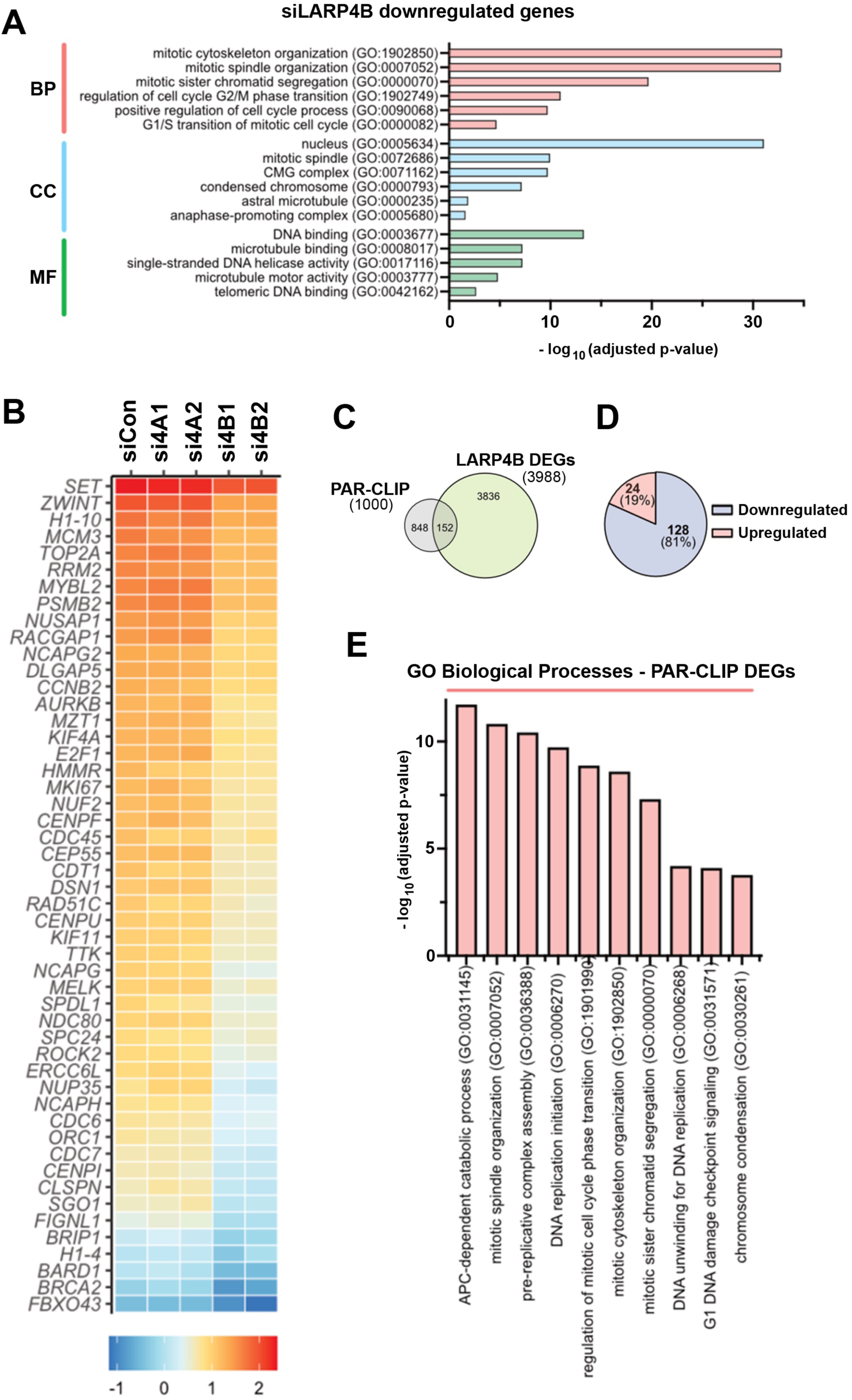
Identification of LARP4A and LARP4B targets. (A) Gene ontology (GO) terms for Biological Process (BP), Cell Component (CC) and Molecular Function (MF) significantly enriched for, following analysis of significantly downregulated genes in LARP4B knockdown PC3 cells compared to siControl cells. (B) Heatmap depicting relative expression (bottom scale Z score) of top 50 mitosis-related genes across siControl (siCon), siLARP4A (si4A1, si4A2) and siLARP4B (si4B1, si4B2) knockdown PC3 cells. (C) Venn diagram showing overlap between mRNAs that feature as differentially expressed genes (DEGs) between LARP4B knockdown and control cells in bulk RNAseq, and candidate direct mRNA targets (top 1000) of LARP4B as shown by PAR-CLIP analysis by Kuspert et al^25^. (D) Pie chart demonstrating the proportion of upregulated and downregulated DEG/PAR-CLIP genes that overlap with bulk RNAseq analysis of LARP4B knockdown PC3 cells compared to controls. (E) Gene ontology (GO) term for Biological Processes (BP) significantly enriched for, following analysis of significantly downregulated putative mRNA target genes of LARP4B, in LARP4B knockdown PC3 cells compared to controls. See also Figure S5.

Cross-examination of mRNA abundance changes from our RNAseq with putative mRNA binding targets of LARP4B, identified in a previous PAR-CLIP study,^25^ revealed a subset of 152 cellular mRNAs that may be regulated directly by LARP4B at the steady-state level (Figure 5C). Of these, more than 80% were downregulated upon LARP4B depletion (128 genes), implying that LARP4B plays a major role in stabilising these mRNA targets (Figure 5D). Notably, several genes with gene ontology (GO) terms for mitosis and cell cycle-related processes were included in this group, specifically cyclins (*CCNB1*, *CCNE2*), cyclin-dependent kinases (*CDK1/2*) and their inhibitors (*CDKN1A/B/C)* and mitotic checkpoint kinases (*AURKA/B*; Figure 5E). We confirmed downregulation of several of the mitosis-related candidate targets of LARP4B, namely *AURKB, CCNB1, CCNE2* and *E2F1* both at the mRNA and protein levels upon siRNA LARP4B depletion (Figures S5A-C). Thus, LARP4B appears to regulate the steady-state level of mRNAs coding for genes involved in cell cycle, in agreement with its role in cell cycle and proliferation, and this function appears to be distinct from LARP4A.

Recent work has reported that LARP4A localises to mitochondria and regulates mitochondrial activity, and this may contribute to the cell proliferation phenotype observed.^30, 55^ However, thus far no data are available for LARP4B. Our cell fractionation revealed that, besides the cytoplasm, LARP4A and LARP4B were expressed at comparable levels in mitochondrial fractions as well as in the nucleus to varying degrees in both MG63 and PC3 cells (Figure 6A). Immunofluorescence and confocal microscopy analyses corroborated this, demonstrating co-localisation of LARP4A and LARP4B with the mitochondrial marker, TOM20 (Figures 6B and S6A). There was a strong and largely equivalent co-localisation of both LARP4A/TOM20 and LARP4B/TOM20 in PC3 cells (Figures S6B-E), whereas in MG63 cells LARP4B showed a higher degree of co-localisation with TOM20, compared to LARP4A (Figures 6C-F). These results may suggest that both LARP4A and LARP4B may affect cell proliferation via regulating mitochondria activity.

**Figure 6.**
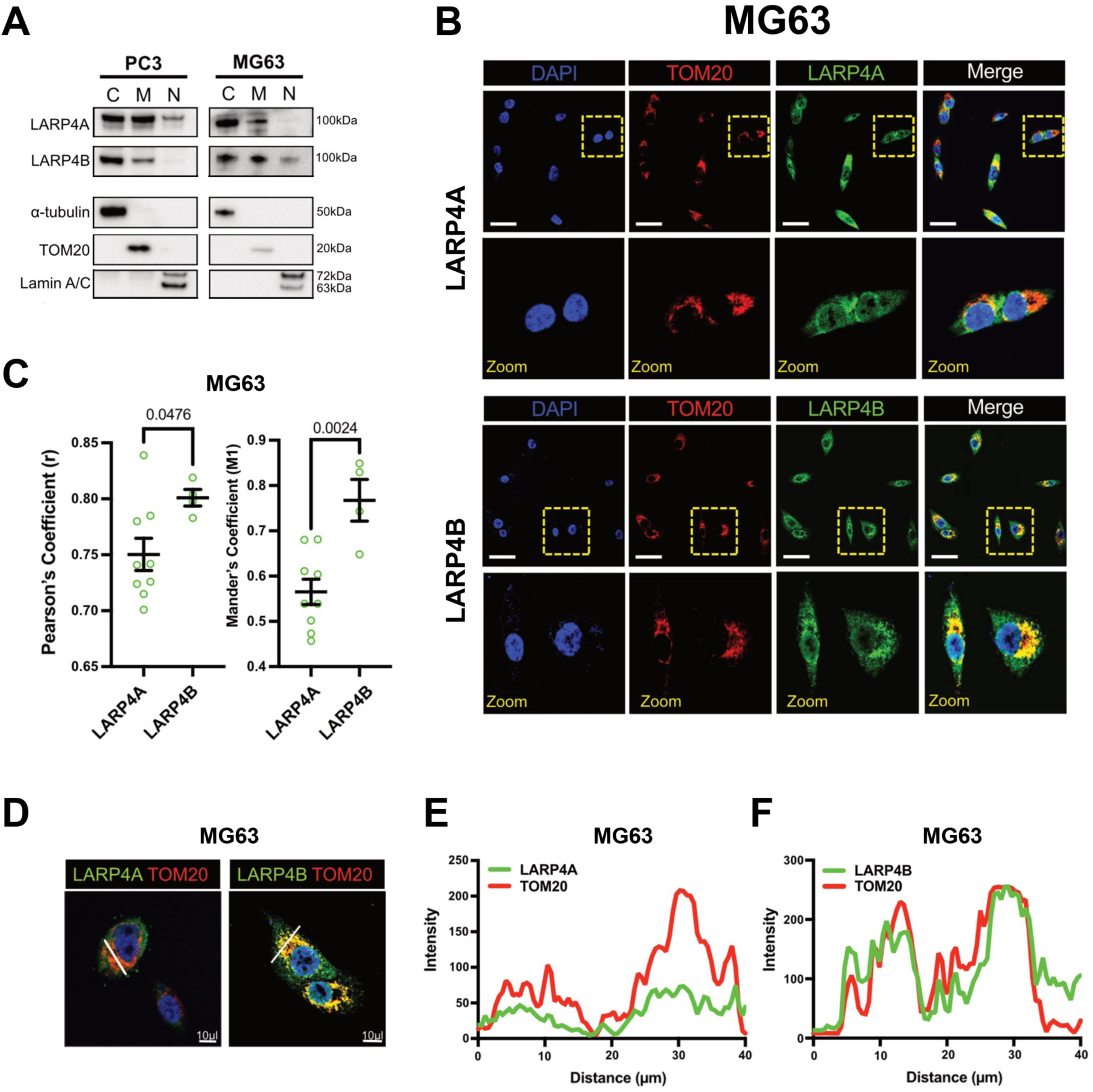
Expression and localisation of LARP4A and LARP4B in mitochondria. (A) Western blot analysis of endogenous LARP4A and LARP4B in PC3 and MG63 cells following fractionation of cytoplasm (C), mitochondria (M) and nuclei (N). α-tubulin, TOM20 and Lamin A/C are used as respective controls to determine fraction purities. (B) Immunofluorescence detection of TOM20 and co-localisation with LARP4A (*top*) and LARP4B (*bottom*) proteins with DAPI counterstain in MG63 cells. Scale bars, 25µm. Areas in yellow squares are depicted at higher magnification (zoom) in lower panels. (C) Pearson’s and Mander’s coefficient analyses showing a higher degree of LARP4B/TOM20 co-localisation compared with LARP4A/TOM20 in MG63 cells. The data represent the mean ± SEM, n=6 from three independent experiments. p-values are indicated. (D-F) Fluorescent intensity line plot profiling demonstrating co-localisation of LARP4A and LARP4B with the TOM20 mitochondrial marker in MG63 cells. White lines in (D) depict the areas used for LARP4A/TOM20 (E) and LARP4B/TOM20 (F) line plot analyses. Scale bars in (D), 10µm. See also Figure S6.

Finally, we investigated whether apoptosis contributed to the reduction in prostate cancer and osteosarcoma cell number upon LARP4A/4B protein knockdown, given the previous suggestion that LARP4B modulates apoptosis in glioma cells.^38^ While a slight increase in the proportion of apoptotic cells was observed in LARP4A- and LARP4B-depleted cells compared to controls, this was not statistically significant (Figures S7A and S7B). Confirming these results, no change in apoptosis was detected in cells overexpressing LARP4A or LARP4B (Figures S7C and S7D). Notably, whilst mRNA levels of the pro-apoptotic factor *BAX* increased upon overexpression of LARP4B in glioma cells,^38^ this was unaltered in LARP4A- or LARP4B-depleted PC3 or MG63 cells (Figures S7E and S7F). Moreover, no changes in mRNA levels were observed in the RNAseq data for other pro-apoptotic (*BAD, PUMA*) and anti-apoptotic factors (*BCL2, BCLXL* and *MCL-1*) (data not shown).

Taken together, loss- and gain-of-function studies suggest that LARP4A and LARP4B promote cancer cell proliferation and survival. This appears to be independent of apoptosis and, primarily for LARP4B, attributed to specific changes in cell cycle and mitosis regulation.

### LARP4A and LARP4B induce morphological changes and promote cancer cell migration *in vitro*

Morphological transformation and increased migration are well-established hallmarks of the transformed phenotype. LARP4A has previously been reported to regulate both cell morphology and migration,^34–36^ but LARP4B has not yet been explored in this context. Depletion of LARP4A or LARP4B in both PC3 and MG63 cells induced increased cell perimeter, elongation and protrusion formation compared to controls (Figures 7A-F). Conversely, overexpression of LARP4A or LARP4B resulted in higher cell circularity and loss of protrusions in both cell types (Figures 7G-L), suggesting that these proteins promote cell rounding. To determine whether the observed cell morphological features may be due to differences in epithelial-to-mesenchymal transition (EMT), the expression of EMT biomarkers was examined. Neither depletion nor overexpression of either LARP4A or LARP4B significantly altered the expression of E-cadherin (PC3 cells) or vimentin (MG63 cells) (Figures S8A and S8B), indicating that these proteins are unlikely acting through EMT. We next determined whether these morphological alterations affected cell migration. LARP4A/4B-depleted PC3 and MG63 cells inhibited cell migration in a scratch assay (Figures 8A-D). To rule out that reduction in wound healing effects may, at least in part, arise from the reduced proliferation rates in LARP4A and LARP4B-depleted cells, we tracked the migratory paths of random, non-proliferating single cells over time. Consistent with previous results, knockdown of LARP4A and LARP4B significantly reduced cell velocity, accumulated and Euclidean distance in both PC3 and MG63 cell lines (Figures 8E-H and Supplementary Videos). Finally, transwell assays demonstrated that silencing of LARP4A, and to a greater extent LARP4B, significantly impaired chemoattractant-mediated cell migration of all cell lines (Figures 8I-L, S9A, and S9B).

**Figure 7.**
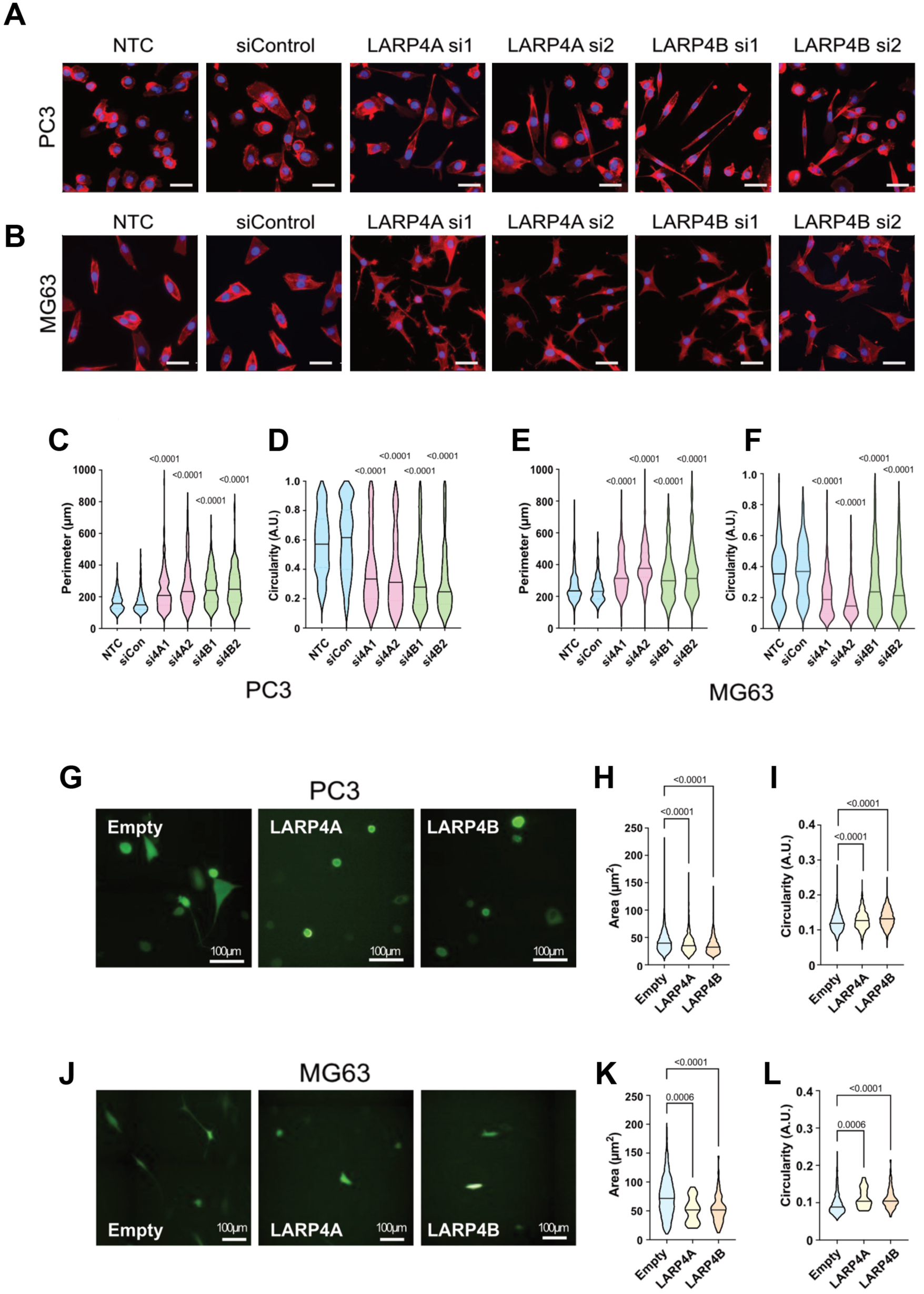
LARP4A and LARP4B regulate cell morphology. (A and B) Immunofluorescence images of PC3 (A) or MG63 (B) control (NTC, siControl) and LARP4A (siLARP4Asi1, siLARP4Asi1) or LARP4B (siLARP4Bsi1, siLARP4Bsi2) knockdown cells stained with phalloidin and DAPI for morphological analyses. Scale bars, 50µm. (C-F) Morphological analysis of PC3 (C and D) and MG63 (E and F) cell perimeter and circularity as indicated. For all violin plots, the line depicts the mean, n=178-220 cells from 3 independent experiments. (G) Immunofluorescence images of PC3 cells overexpressing GFP-LARP4A, GFP-LARP4B or empty GFP plasmid. Scale bars, 100µm. (H and I) Cell area (H) and circularity (I) of PC3 cells overexpressing LARP4A and LARP4B versus an empty plasmid control. Lines in violin plots depict the mean, n=600-1000 cells from 3 independent experiments. (J) Immunofluorescence images of MG63 cells overexpressing GFP-LARP4A, GFP-LARP4B or empty GFP plasmid. Scale bars, 100µm. (K and L) Cell area (K) and circularity (L) of MG63 cells overexpressing LARP4A and LARP4B versus an empty plasmid control. Lines in violin plots depict the mean, n=600-1000 cells from 3 independent experiments. See also Figure S7.

**Figure 8.**
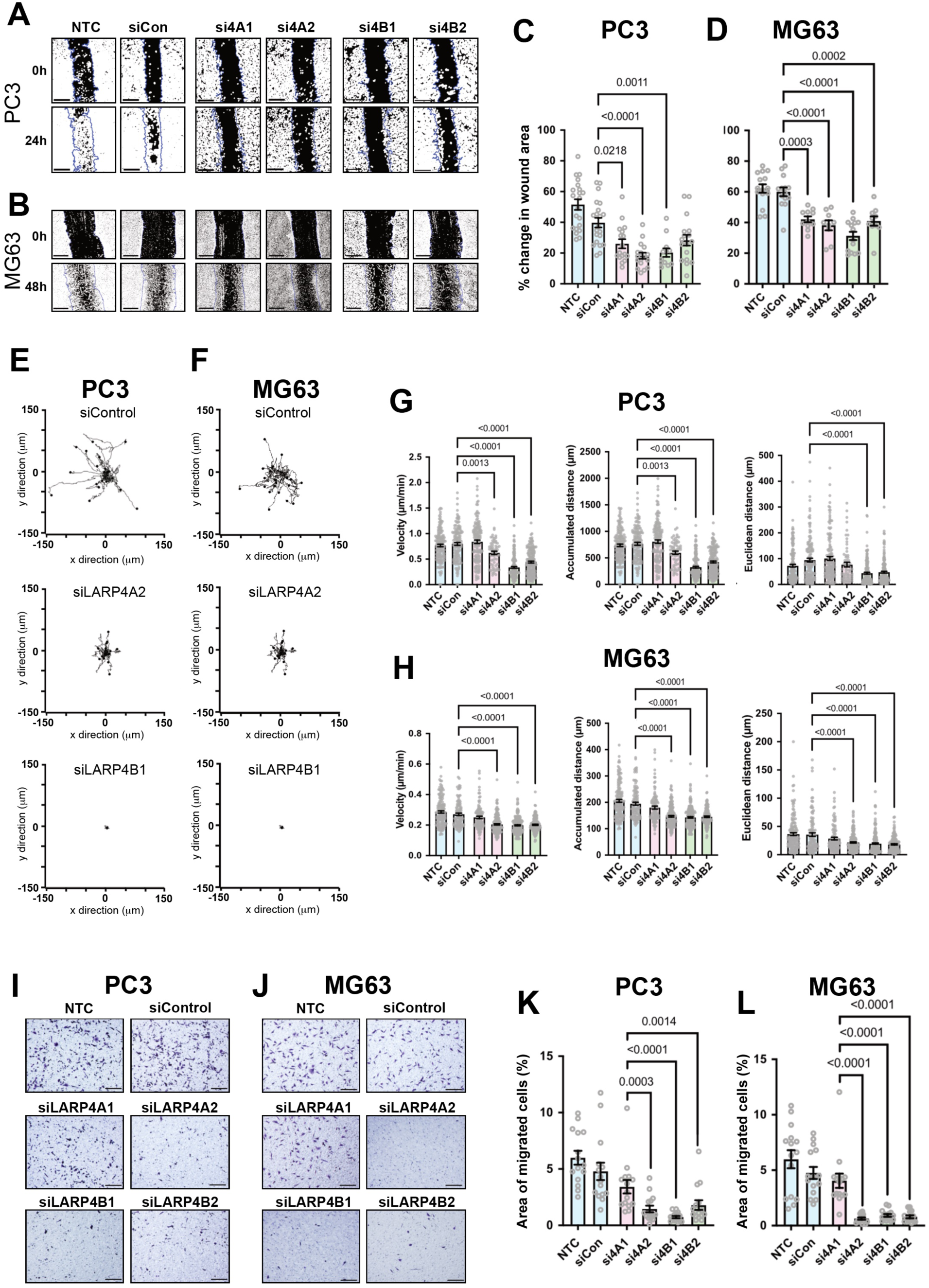
LARP4A and LARP4B regulate cell migration and invasion. (A and B) Wound closure assays in PC3 (A) and MG63 (B) cells over 24-48h either non-transfected (NTC) or transfected with non-targeting control (siCon) and siRNAs targeting LARP4A (si4A1, si4A2) or LARP4B (si4B1, si4B2) as indicated. Blue lines depict the area of wound at t=0h and black/white image shows the mask used for wound area analysis. Scale bars, 200µm. (C and D) Quantification of would area closure by PC3 (C) and MG63 (D) cells over the experimental time course shown in (A and B). The data indicate the mean ± SEM, n=12-22 from 3 independent experiments. p-values are indicated. (E and F) Individual migration tracks of 20 randomly selected PC3 (E) and MG63 (F) control (siControl), siLARP4A or siLARP4B knockdown cell tracks during a 16h time lapse experiment. (G and H) Migration analysis (velocity, accumulated distance and Euclidean distance) following quantification of cell tracks from PC3 (G) and MG63 (H) cell either non-transfected (NTC, siCon) or transfected with siRNAs targeting LARP4A (siA1, siA2) or LARP4B (siB1, siB2). The data represent the mean ± SEM, n=65-180 cells from 3 independent experiments. p-values are indicated. (I and J) Brightfield images of transwells stained with crystal violet following a 24h migration assay of PC3 (I) and MG63 (J) control (siControl), siLARP4A (4A1, 4A2) or siLARP4B (4B1, 4B2) knockdown cells. Scale bars, 200µm. (K and L) Quantification of migrated PC3 (K) and MG63 (L) cells over 24h of either non-transfected cells (NTC, siCon) or cells transfected with siRNAs targeting LARP4A (siA1, siA2) or LARP4B (siB1, siB2). The data represent the mean ± SEM, n=15 from 3 independent experiments. p-values are indicated. See also Figure S9.

Taken together, our results show that both LARP4 paralogs induce morphological changes that are associated with transformed cells and promote cancer cell migration which may underpin their function in promoting the metastatic potential of cancer cells *in vivo*.

## DISCUSSION

The La-related proteins are emerging as key RBPs with essential roles in RNA biology, but their contributions to disease, including cancer, are not well understood. For the paralogs LARP4A and LARP4B, their roles in cancer progression are only just beginning to be recognised. In this work, we demonstrate that LARP4A/4B depletion inhibits osteosarcoma and prostate cancer formation and growth *in vivo*, advocating for pro-tumorigenic roles of these proteins in these cancers. Additionally, we performed a long overdue systematic comparative analysis of these paralogs, identifying similarities and differences between these two proteins and uncovering potential mechanisms underlying their functions.

The most striking observation was the marked inhibition of tumour growth in xenografts lacking LARP4A or LARP4B in human cancer lines of different origins, mesenchymal and epithelial, as represented by osteosarcoma (MNNG/HOS; MG63) and prostate cancer (PC3) cells, respectively. This is consistent with the high endogenous expression of both LARP4 paralogs in osteosarcoma and prostate cancer. Silencing of either LARP4A or LARP4B inhibited cell viability *in vitro* and the mitotic indices of both prostate cancer and osteosarcoma cell xenografts *in vivo*, suggesting that both LARP4 proteins are positive regulators of cell proliferation. The well-established role of LARP4A and LARP4B in sustaining translation rates^23–25^ plausibly impacts on cell proliferation, and this was further corroborated here by our RNAseq analysis reporting elevated mRNA levels of negative regulators of translation and promoters of polyA tail shortening in LARP4A/B depleted cells (*AGO2/4, CPEB* proteins, *BTG2, TNRC6C* and *ZFP36;* data not shown). However, additional cellular mechanisms are likely to play a role. While apoptotic rates were not affected by either LARP4A or LARP4B knockdown, our multiparametric analysis dissecting the consequences of each LARP4 protein knockdown on individual cell cycle stages revealed a novel role for LARP4B in promoting cell cycle progression, specifically affecting entry into mitosis of both cancer cell lines. This was validated by our unbiased genome-wide analyses, revealing that LARP4B regulates a series of targets with GO terms enriched in the regulation of mitosis, spindle organisation and cell cycle phase transition. Cross-examination of differentially expressed genes and candidate LARP4B mRNA targets from a previous PAR-CLIP study^25^ suggests that LARP4B may directly modulate a subset of its RNA targets at the steady-state level to promote mitotic progression. Specific targets that are co-ordinately regulated by LARP4B in a direction that is consistent with mitotic entry and general cell cycle progression include several cyclins and other key cell cycle regulators, namely *AURKA/B, E2F1, CCNB1, CCNE2, CDC7, RCF3, MKI67, BRCA2, CDK1/2* and *CDKN1A/B/C.* Our evidence that LARP4B upregulates these transcripts and enhances translation of *AURKB, E2F1, CCNB1* and *CCNE2* mRNAs establishes these as *bona fide* LARP4B targets with tumour promoting functions. Indeed, for many cancers, including prostate cancer and osteosarcoma, altered cell cycle control represents a hallmark of tumour progression^2^ and cyclins and cell cycle kinases are well-established targets for therapy.^56, 57^ For example, AURKB is a mitotic checkpoint kinase dysregulated in many tumours, being linked to tumour cell invasion, metastasis and drug resistance,^58^ but is also overexpressed in osteosarcoma^59^ and its expression directly correlates with Gleason grade in prostate cancer.^60^ The bidirectional transcriptional regulation of *CCNE2* and *E2F* is critical for cell cycle progression. Moreover, E2F binds to the retinoblastoma (RB1) tumour suppressor protein which is mutated in many cancers and, together with TP53, is an established driver of prostate cancer and represents a major germline mutation in osteosarcoma.^61–63^ Taken together, we have identified a subset of regulated cell cycle, cell proliferation and oncogenesis-related mRNAs that potentially represent a novel LARP4B RNA regulon,^64, 65^ mediating LARP4B’s effects in driving cancer progression, and that this appears to be a non-overlapping function with LARP4A.

It has been suggested recently that LARP4A affects cell proliferation rates by regulating levels of proteins essential to mitochondrial function.^55^ Through its recruitment to mitochondria by the scaffolding factor dAKAP1, LARP4A acts as a general translation factor for the local synthesis of metabolic enzymes that operate the TCA cycle and promote mitochondrial electron transport. LARP4 can also bind to nuclear-encoded mitochondrial mRNAs, regulating their translation.^30, 55^ LARP4A and LARP4B have always been described as cytoplasmic proteins, however, our demonstration that both co-localise to the mitochondria in prostate cancer and osteosarcoma cells supports an essential role for these paralogs in mitochondrial function. To our knowledge, this is the first evidence of mitochondrial localisation of LARP4B and is consistent with the identification of mitochondria-associated genes in the LARP4B interactome.^25^ Together with our bulk RNAseq analysis showing that knockdown of either LARP4A or LARP4B resulted in overexpression of genes enriched for the response to oxidative stress pathways, metabolism and glycolysis (data not shown) our data further endorse the importance of LARP4 proteins in mitochondrial-related activities. While the functional implications of these observations remain to be investigated, a possible crosstalk between energy production and LARP4 protein function at mitochondrial membranes may be consistent with the capacities of LARP4A and LARP4B in energetically demanding processes such as cellular proliferation, migration and cancer development.^66–68^ In addition to subcellular localisation to mitochondria, we demonstrated additional nuclear localisation *in vitro* and significantly, in primary tumours and metastatic lesions *in vivo*, which hints at a more complex functional profile than previously anticipated. Whereas LARP4A was found to shuttle with PABPC1 to the nuclei during accelerated mRNA decay,^69^ this is the first evidence of a nuclear localisation for LARP4B. This might suggest that the roles of LARP4A and LARP4B in RNA regulation extend to nuclear transcripts and/or involve coordinated cytoplasmic/nuclear activities, and notably, LARP4A and LARP4B have both been identified as nuclear RNA interactors in a global capture study of the human cell nucleus.^70^ Investigating further the functional significance of such subcellular localisations, and how these contribute to malignant cell growth and migration would be of prime importance.

Knockdown of LARP4A and LARP4B in osteosarcoma cells markedly reduced their ability to seed and expand in lung tissue in a xenograft metastasis model. This is consistent with the high levels of LARP4 proteins in lung nodules formed in a spontaneous osteosarcoma metastasis model, suggesting for the first time, significant pro-invasion and pro-metastatic roles for these proteins. Whether LARP4 proteins regulate extravasation or have specific interactions with endothelial cells is not yet known; however, our unbiased transcriptomic analysis of cells lacking either LARP4A or LARP4B revealed targets that modulate cell junction assembly, cell migration and ECM organisation and (data not shown). Indeed, our *in vitro* studies demonstrate LARP4A and LARP4B’s regulation of cell morphology, wound healing and migratory capacity, all characteristic features of metastatic cells. Notably, LARP4A had previously been categorised as a cell morphology and migration regulator, a property that is likely to be context-dependent,^35, 36^ whereas to our knowledge this is a new cellular function assigned to LARP4B. Interestingly, the effects of LARP4A and LARP4B, particularly for epithelial-derived PC3 cells, appear to be independent of any changes in genes associated with epithelial-mesenchymal transition (EMT) which is a widely accepted mechanism of carcinoma cell metastasis. Moreover, the emerging role of mitochondrial activity in regulating cancer cell migration and metastasis^71–73^ provides a potential novel regulatory link between LARP4A/B mitochondrial localisation and their biological effects.

In summary, we provide the first comparative *in vivo* study of the functions and carcinogenic signatures of LARP4A and LARP4B in malignancy development and metastasis in specific cancer models and found that their pro-tumorigenic capacities in this context are largely shared. Uncovering targets and mechanisms of regulation for LARP4 proteins indicate that although some cellular functions may have been preserved throughout evolution, mechanistic differences do also exist. Indeed, the increasing investigations of LARP4A and LARP4B in different cancer models indicate complex cell-, tissue- and context-dependent functions of LARP4 proteins in tumour progression. We suggest that LARP4A and LARPB be added to the growing list of RBPs that play important roles in osteosarcoma^74^ and prostate cancer,^75^ and it will be important to address the intricacies of the underlying molecular mechanisms, cell/tissue-specific RNA targets and regulatory networks behind LARP4A and LARP4B activities in future functional studies. Nevertheless, our combined *in vitro* and *in vivo* findings provide unambiguous evidence of the pro-migratory, growth-promoting and oncogenic potentials of LARP4A and LARP4B *in vitro* and *in vivo,* that may be exploited for biotechnological and therapeutic applications.

## METHODS

### Constructs

DNA fragments of LARP4A and LARP4B cDNA were subcloned previously into the pCB6-GFP plasmid to obtain N-terminal GFP-tagged LARP4 proteins. cDNA of LARP4A was obtained through restriction enzyme digest of the pCMV2-FLAG-LARP4A plasmid, kindly donated by Dr Richard Maraia (NIH).^24^ LARP4B cDNA was subcloned similarly from a LARP4B-IMAGE clone (Sources Bioscience). Both cDNA fragments were separately purified and ligated into the empty pCB6-GFP plasmid, and the resulting pCB6-GFP-LARP4A and pCB6-GFP-LARP4B plasmids were verified by sequencing (Sources Bioscience).

### Cell culture

MG63 cells were purchased from ATCC (CRL-1427), PC3 cells were donated by Dr. Magali Williamson (King’s College London, KCL) and MNNG/HOS cells previously transduced with a dual GFP/Luciferase reporter^49^ were kindly donated by Professor Alison Gartland (University of Sheffield). Osteosarcoma cells (MG63, MNNG/HOS) were cultured in α-MEM containing ultra-glutamine (Lonza) and PC3 cells were cultured in RPMI 1640 (Thermofisher) supplemented with 2mM L-Glutamine (Merck). All media were supplemented with 10% FBS (Lonza) and 1% penicillin/streptomycin (Merck) and cultured at 37°C in a humidified incubator (5% CO_2_). Cells were cultured at low passage numbers and passaged consistently using Trypsin-EDTA 0.25% (Merck) and tested regularly for mycoplasma contamination (all cells used for experiments were mycoplasma negative at the time of testing).

### Cell transfections

To overexpress GFP-tagged LARP4A and LARP4B, cells were plated at high density 24h prior to transfection using Lipofectamine 2000 (Thermofisher) according to manufacturer’s instructions. Transient transfections were performed using Lipofectamine and DNA diluted in Opti-MEM serum-free medium (Thermofisher) at a 3:1 ratio. Liposome/DNA complexes were added to cells cultured in antibiotic-free medium and incubated for a further 24h before being used for experiments. An empty plasmid control (pCB6-GFP-empty) was used as a control for overexpression experiments.

To deplete LARP4A and LARP4B, two siRNAs targeting each gene (Horizon Discovery) were transfected into cells plated at high density 24h prior, alongside a non-transfected control (NTC, lipofectamine and no siRNA) and a non-targeting siRNA control (lipofectamine and siControl). Cells were transfected at a final concentration of 10nM siRNA using Lipofectamine RNAiMAX (Thermofisher) according to manufacturer’s instructions. In short, siRNA oligos and Lipofectamine RNAiMAX were diluted and mixed in Opti-MEM serum-free, after which complexes were added to cells in antibiotic-free medium. Transfected cells were then cultured for 72h before harvesting knockdown confirmation and downstream analysis.

### RNA extraction and quantitative PCR

Total RNA was extracted using the RNeasy Mini kit (Qiagen) according to manufacturer’s instructions and quantified using a Nanodrop UV-Vis Spectrophotometer (Thermofisher). RNA-to-cDNA synthesis was performed using dNTPs and mmLVRT reverse transcriptase (Promega) according to manufacturer’s protocol and diluted to a standard concentration for mRNA expression analysis. qPCR was performed using Brilliant II SYBR Green (Agilent) and pre-designed primers

(Integrated DNA Technologies) on the LightCycler 480 II (Roche). Prior to experiments, primer efficiency tests and melt curve analyses were performed to ensure specificity of amplified products from all primer pairs. For relative mRNA expression analysis, Ct values were calculated for each amplified product (performed in triplicate) and normalised β-actin mRNA, which was previously determined to be a suitable housekeeping gene, then further to the experimental control using the 2^-ΛλΛλCt^ method.

### Subcellular fractionation and Western blotting

Cell pellets were lysed using urea lysis buffer (1.6M urea, 4.8% SDS, 19.5mM sucrose). Total protein content was quantified using a bicinchoninic acid (BCA) standard curve assay (Pierce) and reagent absorbance determined using the ClarioStar plate reader (BMG Labtech) at 562nm wavelength. Samples at a standard protein concentration were diluted in Western blot sample buffer consisting of Laemmli buffer (Bio-Rad) and 5% β-mercaptoethanol (Merck). Subcellular fractionation of cytoplasmic, mitochondrial and nuclear fractions were isolated from cells in 10cm dishes using the Standard Cell Fractionation Kit (Abcam) according to manufacturer’s instructions. Fraction purity was determined using SDS-PAGE using the nuclear marker Lamin A/C, the mitochondrial marker TOM20 and α-tubulin as a cytoplasmic protein.

Equal volumes of Western blot samples were loaded onto NuPAGE 4-12% Bis-Tris gels (Thermofisher) alongside molecular weight protein standards (New England Biosciences) and subject to SDS-PAGE in MES running buffer (Thermofisher). Proteins were transferred onto PVDF membranes using a semi-dry transfer system (Bio-Rad) and blocked with 5% (w/v) skimmed milk powder (Applichem) diluted in TBS-Tween (0.1%) at room temperature for 1h. Primary antibodies were diluted in blocking buffer and incubated on membranes overnight at 4°C. After several TBS-Tween washes, membranes were incubated with horseradish peroxidase (HRP)-conjugated secondary antibodies for another hour at room temperature, before the chemiluminescent signal was detected using Clarity ECL substrates (Bio-Rad) on the ChemiDoc Touch imaging system (Bio-Rad). Protein expression was semi-quantified by calculating relative band intensity of target proteins using ImageLab (Bio-Rad) and normalising to those of housekeeping proteins (GAPDH or α-tubulin). Antibodies used: LARP4A (Abcam, ab241489, 1:1000), LARP4B (Abcam, ab197085, 1:2000), GAPDH (Cell Signalling Technology, CST 5174, 1:3000), α-tubulin (CST 2144, 1:3000), E-cadherin (Abcam, ab76055, 1:200), Vimentin (CST 5741, 1:1000), TOM20 (Santa-Cruz, sc-17764, 1:200), Lamin A/C (Santa-Cruz, sc7292, 1:1000), E2F1 (CST 3742, 1:1000), Aurora B (Abcam, ab2254, 1:500), Cyclin B1 (CST, 4138, 1:1000), Cyclin E2 (CST, 4132, 1:1000), Goat anti-rabbit HRP-conjugated IgG (Stratech, 111-035-003, 1:3000), Horse anti-mouse HRP-conjugated IgG (CST, 7076, 1:3000).

### Immunohistochemical and immunofluorescent staining of tissues

Prior to tissue processing, all (bone, tumour and lung) tissues were fixed in 4% PFA overnight and calcified tissues were further decalcified with EDTA. Sections of paraffin-embedded tissues were de-waxed and rehydrated in xylene and a graded ethanol series, respectively, which was then followed antigen retrieval. This was performed in a citrate buffer for soft tissues (lungs, subcutaneous tumours) or Tris-EDTA buffer for bone tissue and osteosarcomas. Sections were blocked for 1h at room temperature in blocking buffer (composed of serum, BSA and Tween-20 diluted in PBS) and primary antibodies, also diluted in blocking buffer, were incubated on tissues overnight at 4°C in a humidified slide box. For immunofluorescence stains, incubation with fluorophore-conjugated secondary antibodies was performed for a further 1h at room temperature. Sections were then washed and mounded with coverslips in Fluoroshield with DAPI (Abcam). For immunohistochemical stains, 3% hydrogen peroxide (Merck) was applied to the sections prior to incubation with HRP-conjugated secondary antibodies for 1h at room temperature. Peroxidase substrates were stained using a DAB kit (VectorLabs), followed by Mayer’s Haematoxylin for 5min. Samples were dehydrated through an increasing ethanol series before being mounted with coverslips using DPX (Merck). For histological staining, sections were stained in Mayer’s Haematoxylin for 8-10min following the same rehydrating graded ethanol series. Following rinsing with dH_2_O and 95% ethanol, an Eosin Y counterstain was employed for a duration between 30 seconds to 2min depending on the tissue type. Samples were rinsed and rehydrated through an ethanol series before being mounted with DPX. Primary antibodies used: anti-LARP4A (Merck, HPA039306, 1:50), anti-LARP4B (Merck, HPA036566, 1:100), anti-Ki67 (Abcam, ab15580, 1:100), anti-GFP (Abcam, ab13970, 1:300), anti-Cleaved Caspase-3 (CST, 9661, 1:200), anti-pHH3(S10) (CST, 9701, 1:400). Secondary antibodies used: Goat anti-Rabbit AlexaFluor 488 (Thermofisher, A11070, 1:200), Goat anti-Rabbit AlexaFluor 546 (Thermofisher, A11035, 1:200), Goat anti-Chick AlexaFluor 647 (Abcam, ab150171, 1:200), Goat anti-Rabbit HRP-conjugated IgG (Stratech, 111-035-003, 1:200). Immunofluorescent lungs from tail vein experiments were imaged using the Axio Observer7 microscope with Apotome 2 (Zeiss). All other histological, immunohistochemical and immunofluorescent tissue stains were imaged using the Hamamatsu 2-HT slide scanner (Hamamatsu). Expression analysis of subcutaneous tumours was performed using QuPath platform for bioimage analysis using the cell detection tools (v.3.0)^76^.

### Immunocytochemistry

Cells were plated at low density following incubation with DNA or siRNA 24h prior to fixation with 4% paraformaldehyde (PFA, Merck) for 15min at room temperature. Permeabilisation was performed using 0.5% Triton X-100 (Merck) for 5min at 4°C and cells were further blocked in 5% goat serum in PBS for 1h room temperature. Cells were incubated in primary antibodies diluted in blocking buffer overnight at 4°C before fluorescent secondary antibody staining in the dark for 1h at room temperature and DAPI counterstaining for 5min prior to imaging. For confocal microscopy, cells which were previously plated on glass coverslips were mounted on slides using ProLong Gold Antifade Mountant (Thermofisher) and left to cure overnight before microscopy on the SP5 microscope (Leica) equipped with 405 Blue Diode, Argon (450-520mm), DPSS 561 and HeNe 633 lasers. For high-content analysis and other non-confocal microscopy, cells were imaged within their plates in PBS using the Operetta CLX high-content analysis system (PerkinElmer) and the Axio Observer7 with Apotome (Zeiss), respectively. Primary antibodies used: LARP4A (Merck, HPA039306, 1:100), LARP4B (Merck, HPA036566, 1:150), TOM20 (Santa Cruz, sc-17724, 1:200). Secondary antibodies/stains: Phalloidin-TRITC (Merck, P1951, 1:400), Anti-rabbit AlexaFluor 488 (Thermofisher, A11070, 1:1000), Anti-rabbit AlexaFluor 546 (Thermofisher, A11035, 1:500), Anti-Mouse AlexaFluor 647 (Thermofisher, A21236, 1:1000).

### Co-localisation analyses

Fixed cells were stained immunocytochemically as described and imaged using confocal microscopy. Pearson’s and Mander’s pixel correlation and co-localisation coefficients were generated using the ImageJ (NIH) “Just Another Co-Localisation Plugin” (JACoP, v.2.0) ^77, 78^ and line density profiles of pixel co-occurrence were calculated using the built-in line and plot profile tool.

### High content multiparametric cell cycle analysis

Multiparametric cell cycle analysis was performed using a previously described technique.^50^ Cells grown in a 96 well plate were incubated with 10µM EdU from the AlexaFluor 647 Click-iT EdU Proliferation kit (Thermofisher) for 8h before being fixed in 4% PFA for 15min at room temperature. Washed cells were permeabilised in 0.5% Triton X-100 before 50µl Click-iT reaction cocktail (mixed according to manufacturer’s instructions) was added to each well for 30min at room temperature. Cells were blocked in 5% goat serum for 1h then incubated with primary pHH3(S10) antibody (CST, 53348, 1:800) overnight at 4°C. Cells were finally incubated with secondary antibody (Thermofisher, A11035, 1:500) and Hoechst (1µg/ml) for 1h at room temperature before washing with PBS and analysing on the Operetta CLS (PerkinElmer). Cells were imaged at 10X using the following channels: Hoechst 33342, AF546, AF647, EGFP (if applicable) using pipeline building blocks as previously described.^50^ Complete cells were initially selected based on the Hoechst channel and the ‘Remove Border Objects’ tool, and singlets/doublets were determined using the PhenoLOGIC machine learning software utilising nuclei property parameters (STAR method). The intensity properties of all channels were calculated for singlets only and used for analysis. As described previously,^50^ frequency distributions of Hoechst intensities of all cells were plotted to find the G1 peak and single-cell Hoechst intensities were normalised to the value of this peak to obtain ‘Normalised DNA content’. This allowed categorisation of cells into cell cycle stages (sub-G1, G1/S and G2/M) based on their normalised DNA content. Cells could be more specifically categorised into S phase using their Hoechst intensities in combination with their mean EdU incorporation, and into M phase with their pHH3 expression. To perform cell cycle analyses on only GFP-LARP4A/LARP4B expressing singlets in overexpression experiments, an adapted pipeline with an additional building block was used to determine GFP^+^ cells using GFP intensity properties.

### Growth assays

For viability assays, CellTitre 96 AQ_ueous_ One Solution MTS (Promega) was added to cells plated at low density according to manufacturer’s instructions and incubated in darkness in a humidified incubator for 3h at 37°C, 5% CO_2_. Absorbances of media containing MTS were then measured at 490nm using the ClarioStar plate reader (BMG LabTech). Each cell condition was measured every 24h over the course of 4d. For cell growth assays, cells were plated in low confluency and left to grow in normal conditions for 72h, after which they were trypsinised and counted using a haemocytometer to determine total cell number.

### Apoptosis assays

Cells were collected following transfection incubations and resuspended in Annexin V binding buffer and Annexin V AlexaFluore 350 conjugate (Thermofisher) or Propridium Iodide (PI, Thermofisher) according to instructions. Cell suspensions were incubated in the dark at room temperature for 15min before being analysed by flow cytometry on the LSRFortessa system (BD Biosciences), alongside positive and negative controls, as well as PI- or Annexin V-only controls to calculate compensation. Thresholding was performed using the DAPI (Annexin V) and PE-Texas Red (PI) channels, as well as the EGFP channel for overexpression experiments to determine GFP^+^ populations. Analysis of apoptotic populations was performed on FlowJo software (BD Biosciences).

### Morphological analysis

Analysis of cellular morphology was performed as previously described.^36^ Cells plated in tissue culture plates pre-coated with Matrigel were fixed and stained using immunocytochemistry as described above. Cells stained with Phalloidin and DAPI were imaged at 10X using TRITC, DAPI and EGFP channels (if applicable, for overexpression experiments). Six regions of interest were taken from each well, with conditions performed in duplicate over three independent experiments. Image files were analysed using ImageJ (NIH). Cell shape parameters, including perimeter, area and circularity, were determined for each cell within the field of view based on phalloidin stains, and calculations for circularity were made (below), producing arbitrary values where 0 indicates a perfect circle and 1 indicates a straight line.

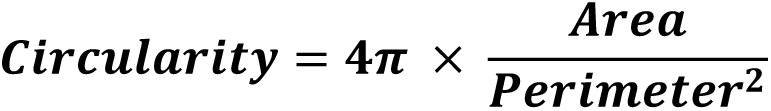

### Migration assays

Determining the influence of LARP4 proteins on cell migration was determined using three independent methods. For scratch assays, cells were plated as a confluent monolayer and serum-starved for 24h prior to the start of the experiment to limit cell division. Wounds were formed in the wells, washed twice and imaged at 4X magnification using phase-contrast time lapse microscopy using the IX81 inverted live-cell light microscope (Olympus) over 24h (PC3 cells) or 48h (MG63 cells). Four regions of interest per scratch were measured over the experiment duration with a frame acquisition of one per 30min. Wound closure over time was quantified using ImageJ, where wound area was quantified at 0h (t=0), and several other time points until 24 or 48h. Wound closure was finally calculated as the percentage change in wound area over time. For single cell migration experiments, cells were plated at low density prior to the start of the experiment in phenol red-free medium. Imaging was performed over 12-18h, with cells being imaged using phase-contrast time lapse microscopy using the IX81 inverted live cell light microscope (Olympus) equipped with MetaMorph time lapse software (Molecular Devices). One frame was captured every 5min and compiled into stack files using MetaMorph software. Stacks were imported into ImageJ (NIH) and cells were manually tracked using the ‘Manual Tracking’ plugin. Cells that divided, died or moved out of the image frame during the course of the experiment were not tracked. Individual cell velocity, accumulated and Euclidean distance were calculated using cell coordinates for each frame using the Chemotaxis and Migration Tool (Ibidi). For transwell migration, cells were seeded onto 8µm pore Transwell chambers (Nunc) containing serum-free medium, which were further placed into wells containing medium supplemented with 10% FBS. The plates were incubated at 37°C, 5% CO_2_, for 24h before transwells were washed, fixed in 4% PFA, permeabilised in methanol and stained with crystal violet for visualisation of migrated cells. Non-migrated cells were removed using a swab and transwell membranes imaged using an Olympus-LS brightfield Stereo Microscope (Olympus) and crystal violet positive cells that had migrated to the other side of the transwell were quantified using ImageJ.

### Ethical approvals and animal usage

All procedures complied with the UK Animals (Scientific Procedures) Act 1986 and were reviewed and approved by the local Research Ethics Committee at KCL. Mice used for subcutaneous and intravital experiments were maintained at the King’s College London (KCL) Biological Services Unit (BSU) under a UK Home Office Project License (P3EBB2E64), while mice used in paratibial xenograft models were maintained at The University of Sheffield (TUoS) BSU Unit under a UK Home Office Project License (PP1668508). For subcutaneous and intravital xenografts, 6-week-old immunocompromised male SCID mice (*CB17/lcr-Prkdc^scid^/lcrlcoCrl*) were used. For paratibial models, 6-week-old female Balb/c nude mice were used. All mice for *in vivo* experiments were purchased from Charles River Laboratories International Inc. and were acclimatised for one week at a minimum in their respective units prior to experimental manipulation. KCL and UoS BSU staff undertook daily husbandry procedures during the experimental courses.

### Xenograft experiments

For subcutaneous models of osteosarcoma and prostate cancer cell xenografts, 1 million viable MNNG/HOS and PC3 cells that had been transfected with control or LARP4A/LARP4B-targeting siRNAs 72h prior (as described above) suspended in a 100µl volume of 1:1 mix of Matrigel:PBS were injected subcutaneously into both flanks of each mouse (2 sites per mouse; 5-8 mice per group, n=15-24). All mice were randomly allocated and experiments were blinded. Mice were monitored for 15d for tumour growth, at which point mice were sacrificed and tumours excised for further analysis. Tumour dimensions were measured using electronic calipers and volume was calculated using the following formula:

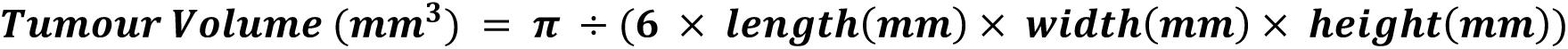

For paratibial models of osteosarcoma xenografts, 500,000 viable MNNG/HOS cells previously transfected with targeting/non-targeting siRNAs were injected onto the surface of both tibiae of anaesthetised mice (isoflurane inhalation, 8 per group, n=24). All mice were randomly allocated and experiments were blinded. Luciferase measurements were taken every other day beginning from day 2 (the day after the injection) – mice were anaesthetised (isoflurane inhalation) and subcutaneously injected with Xenolight D-luciferin substrate (PerkinElmer). Mice were imaged after 5min using the IVIS Lumina II (Caliper Life Sciences), images acquired and luciferase luminescence quantified (total photon flux) using IVIS Living Image software (PerkinElmer). Tumours of control mice became palpable at day 9, after which tibial volumes of mice were measured every 2-3d. Two perpendicular measurements across the tibia were taken and total tibial/tumour volume was calculated using the following formula (M_L_ = Largest measurement; M_S_ = Smallest measurement)

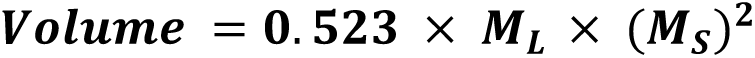

After 14d, mice were sacrificed and tibiae and lungs were dissected for further analysis. Tibiae were fixed for 48h in formalin then switched to 70% ethanol before scanning on a SkyScan 1172 microCT machine (Bruker) at 8µm resolution. The X-Ray source functioned at 200µA, 50kV using a 0.5mm aluminium filter with images captured each 0.7° angle and images were lastly reconstructed using the SkyScan NRecon software (Bruker, v.1.6.9).

### Intravital models of metastasis

For the short term intravital model of metastasis, 1 million viable CMFDA-labelled MG63 cells previously transfected with targeting or non-targeting siRNAs were injected into the tail veins of mice (6 per group, n=18) in 100µl PBS and sacrificed 24h later. For the longer-term intravital model, 5 million viable MNNG/HOS cells previously transfected with targeting or non-targeting siRNAs were injected into mice tail veins in 100µl PBS and mice were sacrificed after 7d. For both models, murine lungs were dissected and fixed for histology and immunofluorescent staining.

### Tissue mRNA expression analyses (TCGA TARGET GTEX)

#### The Cancer Genome Atlas mRNA Expression Database Analysis

Normal tissue versus tumour tissue *LARP4A* and *LARP4B* gene expression analysis was performed using the University of California Santa Cruz (UCSC) Xena browser.^79–81^, mRNA expression data from The Cancer Genome Atlas (TCGA) Therapeutically Applicable Research to Generate Effective Treatments (TARGET) and Genotype Tissue Expression Project (GTEx) study were used. mRNA expression values of *LARP4A* and *LARP4B* were available in normal and tumour tissue across 23 tissue types. Differences between means of normal versus tissue mRNA expression were analysed using a Welch’s t-test in GraphPad Prism (v.9.1.1).

### Bulk RNAseq

RNA was extracted according to the procedure described above from cells previously transfected with siControl siRNA or two siRNAs targeting LARP4A or LARP4B (two independent replicates per condition). High-quality RNA samples were submitted to Genewiz (Bahnofstrasse), which further confirmed high RNA integrity using a BioAnalyzer 2100 (Agilent). Following poly(A) selection and library preparation, mRNA was sequenced using the NovaSeq 6000 system (Illumina; paired end, 150bp). FastQC^82^ confirmed high per base sequence quality and high per sequence quality scores for each individual fastq file and assessed duplicate reads. HISAT2^83^ then mapped reads to the human reference genome (Hg38) and individual count files were generated using ‘featureCounts’.^84^ EdgeR^85, 86^ was finally used for gene normalisation and differential expression testing between groups. The following were group comparisons used: MG63 siLARP4A *or* siLARP4B vs siControl; PC3 siLARP4A *or* siLARP4B vs siControl. Differentially expressed genes (DEGs) were determined using the following standard parameters: false discovery rate (FDR) ≤ 0.05, logFC value of >0.5 (upregulated vs siControl) or <0.5 (downregulated vs siControl). Enrichr^87–89^ was used for gene enrichment analysis using the Gene Ontology (GO) database.

### Statistical analysis

With the exception of RNAseq analysis, which is described in detail above, all statistical analyses were accomplished using GraphPad prism (v.9.1.1). Testing for statistically significant differences in means between two groups was performed using Welch’s t-tests (non-paired), where testing for differences between more than two groups was performed using one-way Analyses of Variances (ANOVA) followed up with Tukey’s multiple comparisons *post-hoc* tests. Statistical significance was defined using the standard threshold of p<0.05. All experiments were performed using a minimum of three independent biological replicates unless otherwise stated, with between two to four technical replicates per sample.

### Materials availability

Any materials generated from this study are available upon request from the corresponding authors/lead contact.

### Data availability

All raw bulk RNAseq data are available in GEO under the accession number GSE222467.

## ACKNOWLEDGEMENTS

The authors thank Prof. Peter Parker (Francis Crick Institute, UK), and Dr. Eric Rahrmann (University of Cambridge, UK) for insightful discussion and critical reading of the manuscript. We thank Drs. Richard Maraia and Sandy Mattijssen (NIH, USA) for the gift of the pCMV2-FLAG-LARP4A plasmid and useful discussions. We are grateful to Dr. Isabel Cruz-Gallardo and Jiazhen Shen for cloning pCB6-GFP-LARP4A and LARP4B plasmids. We thank Dr. Matthias Krause (King’s College London) for providing the IX81 microscope set up for the migration experiments, Dr. Anita Grigoriadis (King’s College London) for help with the RNAseq experiment, Dr. Magali Williamson (King’s College London) for providing us with the PC3 cell line, Prof. Tony Ng (King’s College London) for enabling the *in vivo* studies, and Ms Dhivya Chandrasekaran (King’s College London) for technical help with the xenograft experiments. We thank the BRC Flow Cytometry Core and Biological Services Unit at King’s College London.

This work was supported by a grant from the UK Medical Research Council (MR/N013700/1) awarded to J. Coleman as part of a Doctoral Training Partnership scholarship at King’s College London.

## AUTHOR CONTRIBUTIONS

M.R.C., A.R. and A.E.G. conceived the study. J.C., M.R.C and A.E.G. planned and designed experiments. J.C. L.T., L.K., H.Y., S.H., P.J., A.C., C.C., M.R.C. and A.E.G., performed experiments. V.Y. generated, analysed and provided expert consultation on bulk RNAseq analysis. L.T. and A.G. performed the paratibial xenografts and subsequent analysis. M.R.C. and A.E.G. supervised the study. J.C., M.R.C. and A.E.G. wrote and edited the manuscript.

## FIGURE LEGENDS

## SUPPLEMENTARY FIGURE LEGENDS

**Figure S1.**
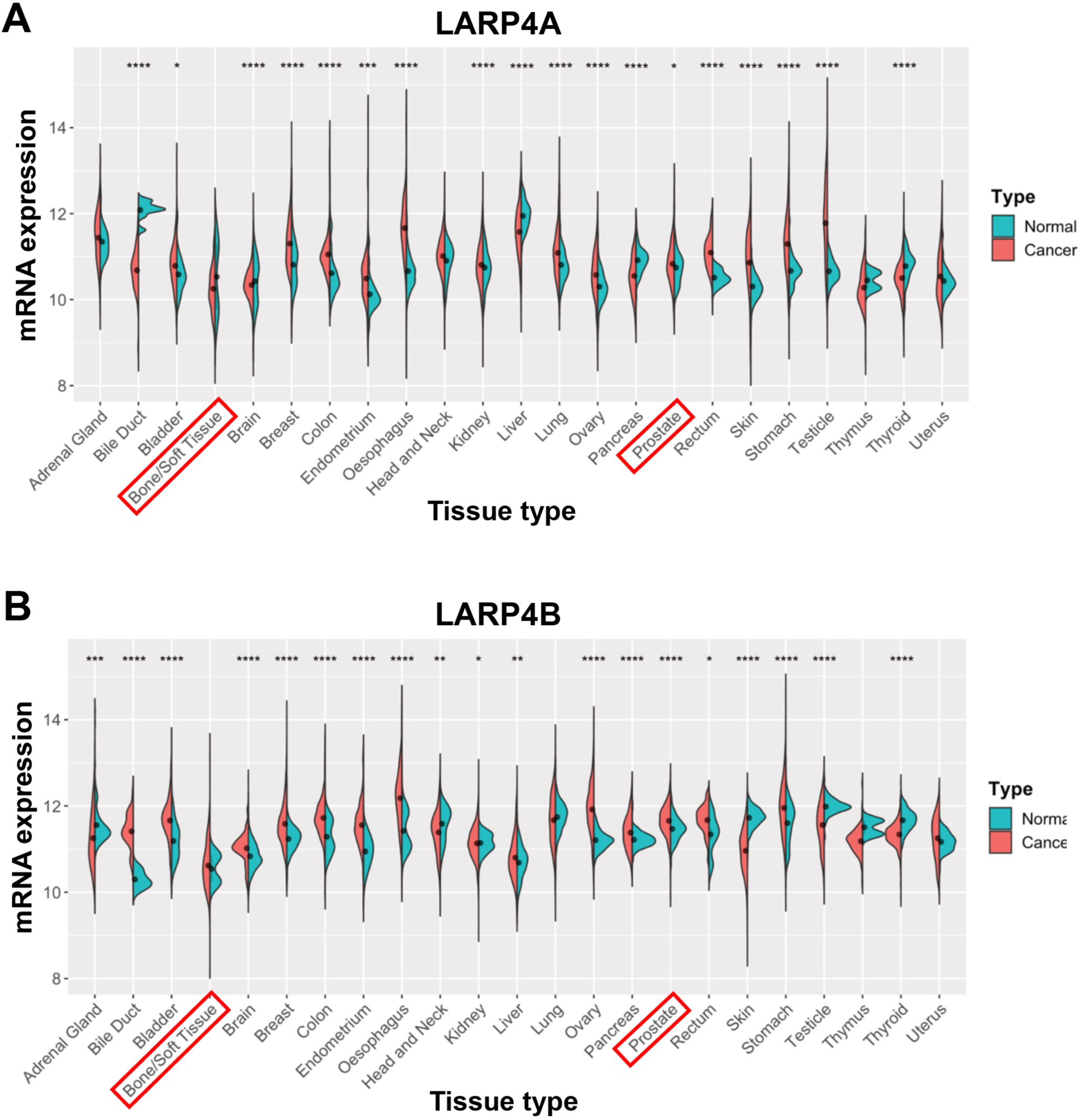
LARP4A and LARP4B expression in different cancers. (A and B) Analysis of *LARP4A* (A) and *LARP4B* (B) mRNA expression across 22 cancer types. Raw RNAseq data (normalised, RSEM counts) from TCGA TARGET GTEx study, obtained from USCS Xena browser (https://xena.ucsc.edu). Sample numbers vary between tissue type, ranging from n=2-1555. Bone and prostate tissues are indicated. The statistical significance of differential expression in normal versus cancer tissue in each tissue is shown. *p<0.05, **p<0.01, ***p<0.001, ****p<0.0001.

**Figure S2.**
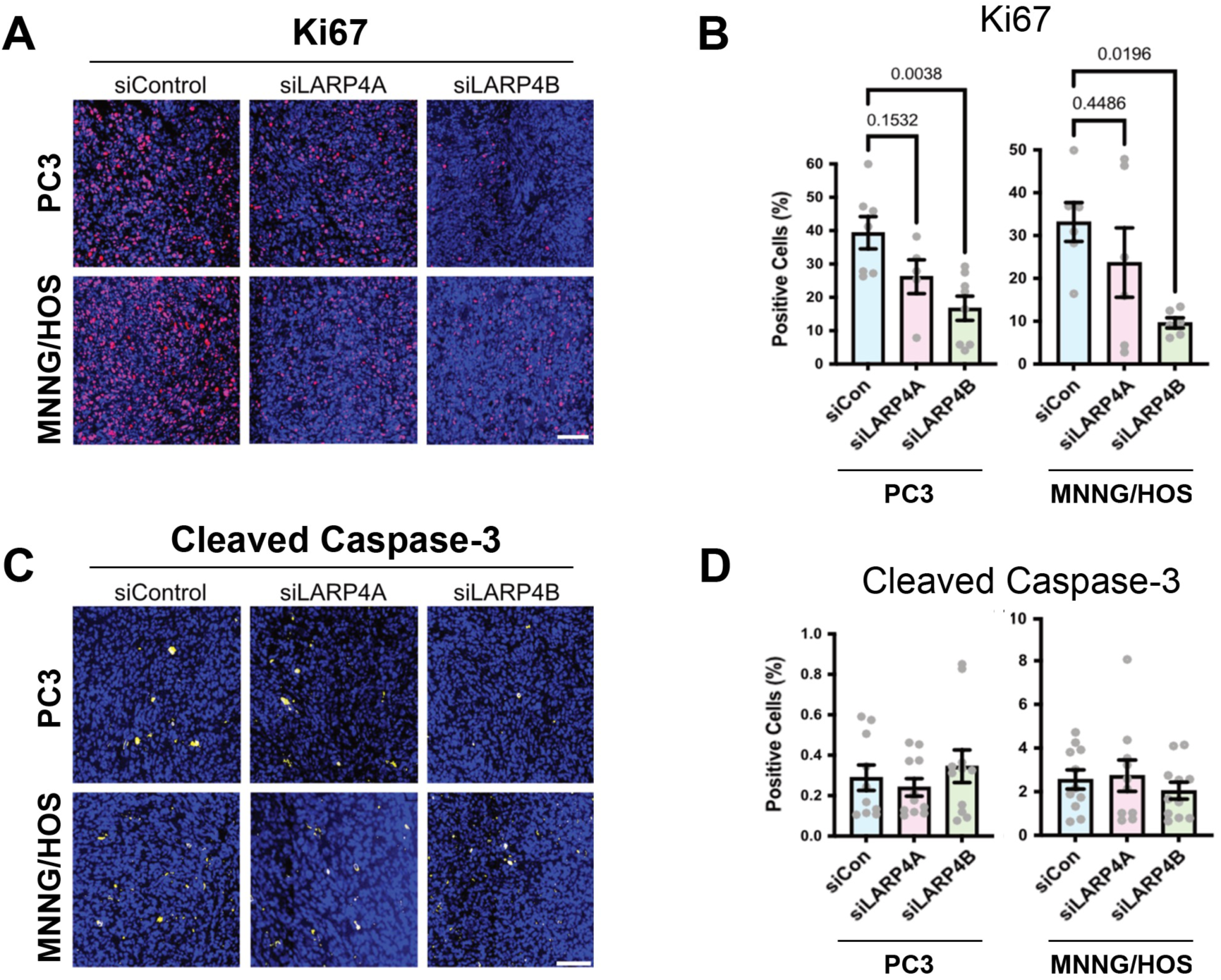
Inhibition of Ki67 and Cleaved Caspase-3 expression following silencing of LARP4A and LARP4B in PC3 and MNNG/HOS cells. (A) Representative immunofluorescence images of Ki67 staining in subcutaneous tumours formed by engraftment of PC3 (*top*) or MNNG/HOS cells (*bottom*) transfected with non-targeting siRNA controls (siControl) or siRNAs targeting LARP4A or LARP4B. Scale bar, 100µm. (B) Quantification of the proportion of cells within PC3 (*left*) and MNNG/HOS (*right*) xenografts expressing Ki67 shown in (A), as a percentage of the total number of cells (DAPI). Bars represent the mean ± SEM per tumour, n=9-16 individual tumours from 6-8 mice. p-values are indicated. (C) Representative immunofluorescence images of Cleaved Caspase-3 staining in subcutaneous tumours formed by engraftment of PC3 (*top*) or MNNG/HOS cells (*bottom*) transfected with non-targeting siRNA controls (siControl) or siRNAs targeting LARP4A or LARP4B. Scale bar, 100µm. (D) Quantification of the proportion of cells within PC3 (*left*) and MNNG/HOS (*right*) xenografts expressing Cleaved Caspase-3 shown in (C), as a percentage of the total number of cells (DAPI). Bars represent the mean ± SEM per tumour, n=9-16 individual tumours from 6-8 mice. p-values are indicated.

**Figure S3.**
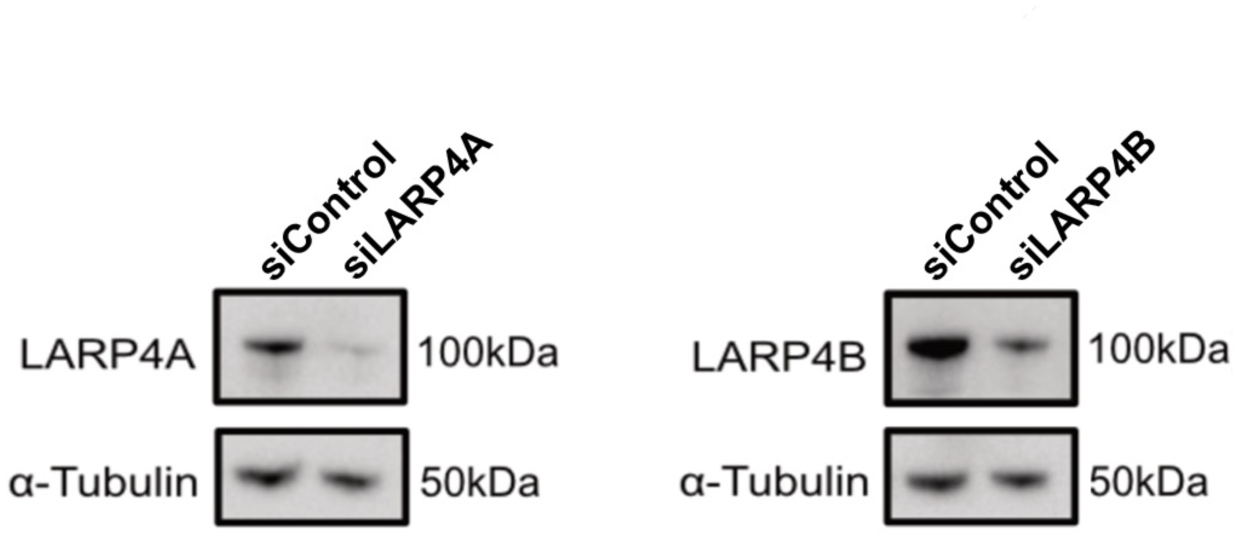
Western blot analysis confirming knockdown of LARP4A and LARP4B expression in MNNG/HOS cells prior to intravenous injection into the tail vein. α-tubulin was used as a loading control.

**Figure S4.**
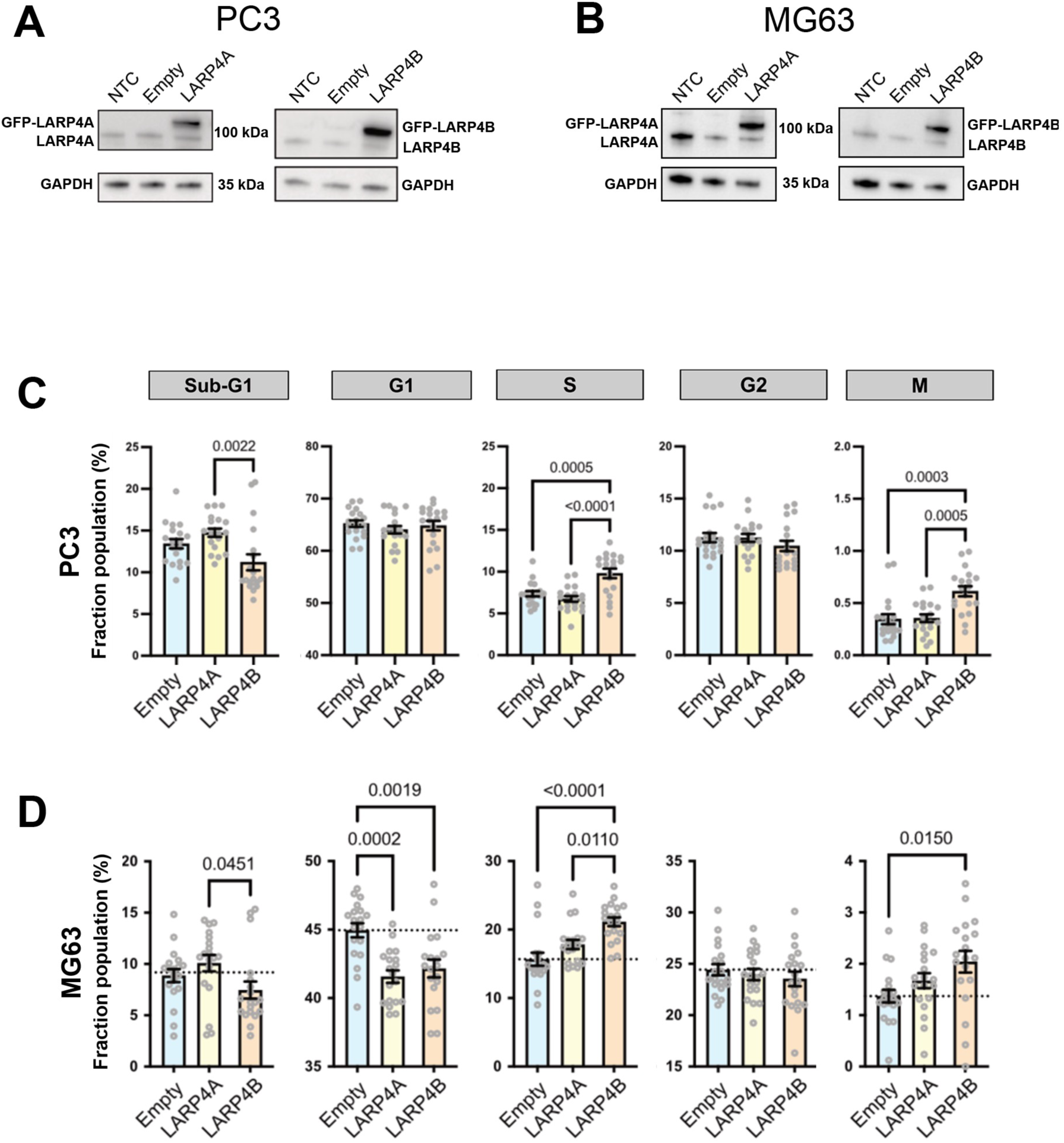
Overexpression of LARP4A and LARP4B regulates cell cycle in PC3 and MG63 cells. (A and B) Western blot analysis of LARP4A and LARP4B overexpression in PC3 (A) and MG63 (B) cells following transfection with GFP-LARP4A or GFP-LARP4B compared to empty non-transfected controls (NTC). GAPDH is used as a loading control. (C and D) Multiparametric cell cycle analysis of PC3 (C) and MG63 (D) cells overexpressing LARP4A, LARP4B or empty plasmid control as indicated. The fraction of the population at each cell cycle stage (sub-G1, G1, S, G2 and M phases) for each cell type is depicted. The data represent the mean ± SEM, n=20 from four independent experiments. p-values are indicated.

**Figure S5.**
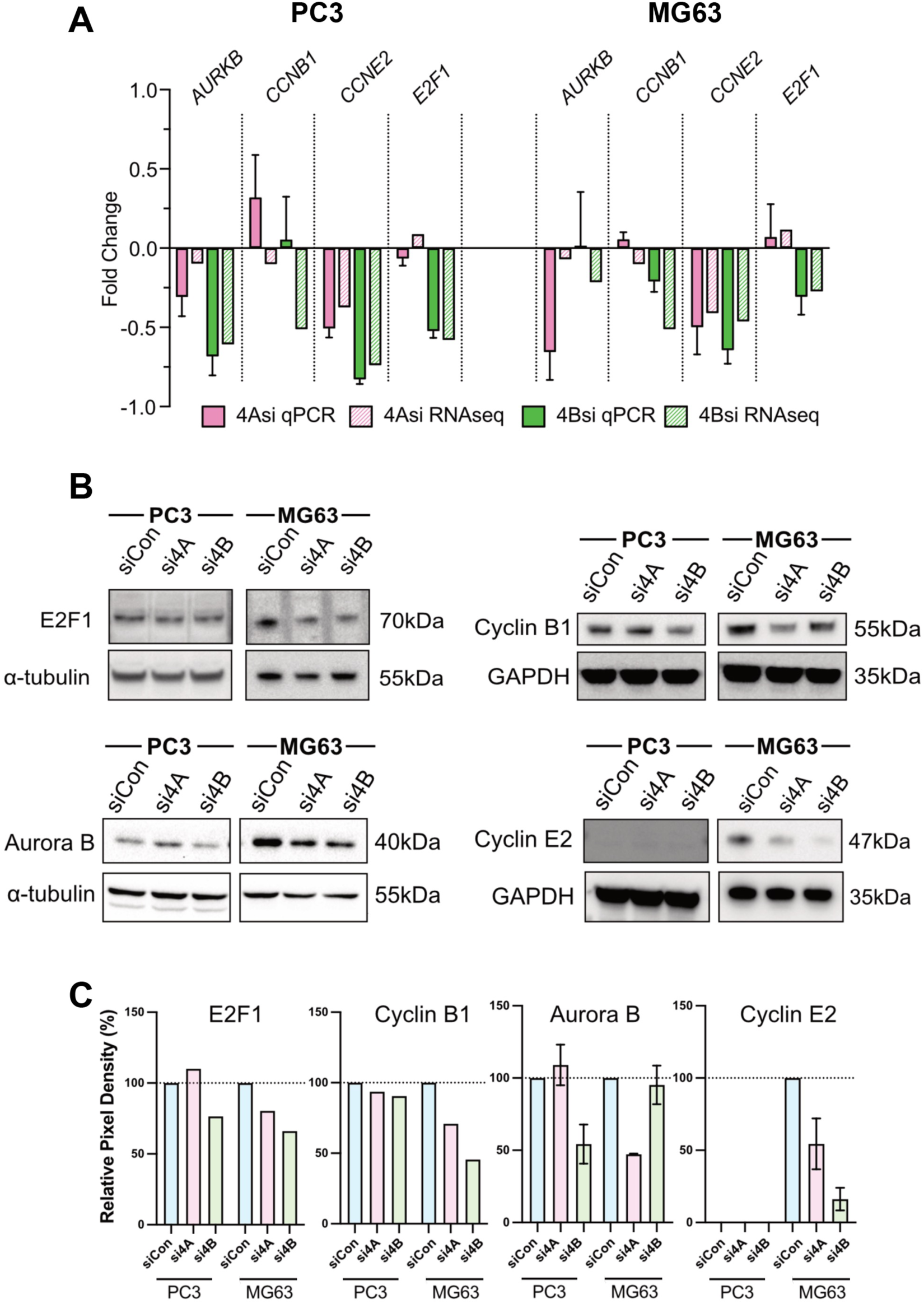
Inhibition of cell cycle targets following silencing of LARP4A and LARP4B in PC3 and MG63 cells. (A) RNA expression of mitotic-related genes detected via RNAseq and validated using qPCR. Expression is shown as fold change expression of LARP4A knockdown (si4A) or LARP4B knockdown (si4B) compared to siRNA controls, from 2-ΛλΛλCt values in qPCR and FPKM in RNAseq. The data represent the mean ± SEM, n=4 (qPCR) or n=2 (RNAseq) per condition. (B) Western blot analysis of E2F1, Cyclin B1, Aurora B and Cyclin E2 protein expression in control (siCon), LARP4A (si4A) and LARP4B (si4B) knockdown PC3 and MG63 cells. α-tubulin or GAPDH are used as loading controls. (C) Quantification of protein expression by densitometry analysis in immunoblots shown in (B), normalised to pixel intensity of respective controls.

**Figure S6.**
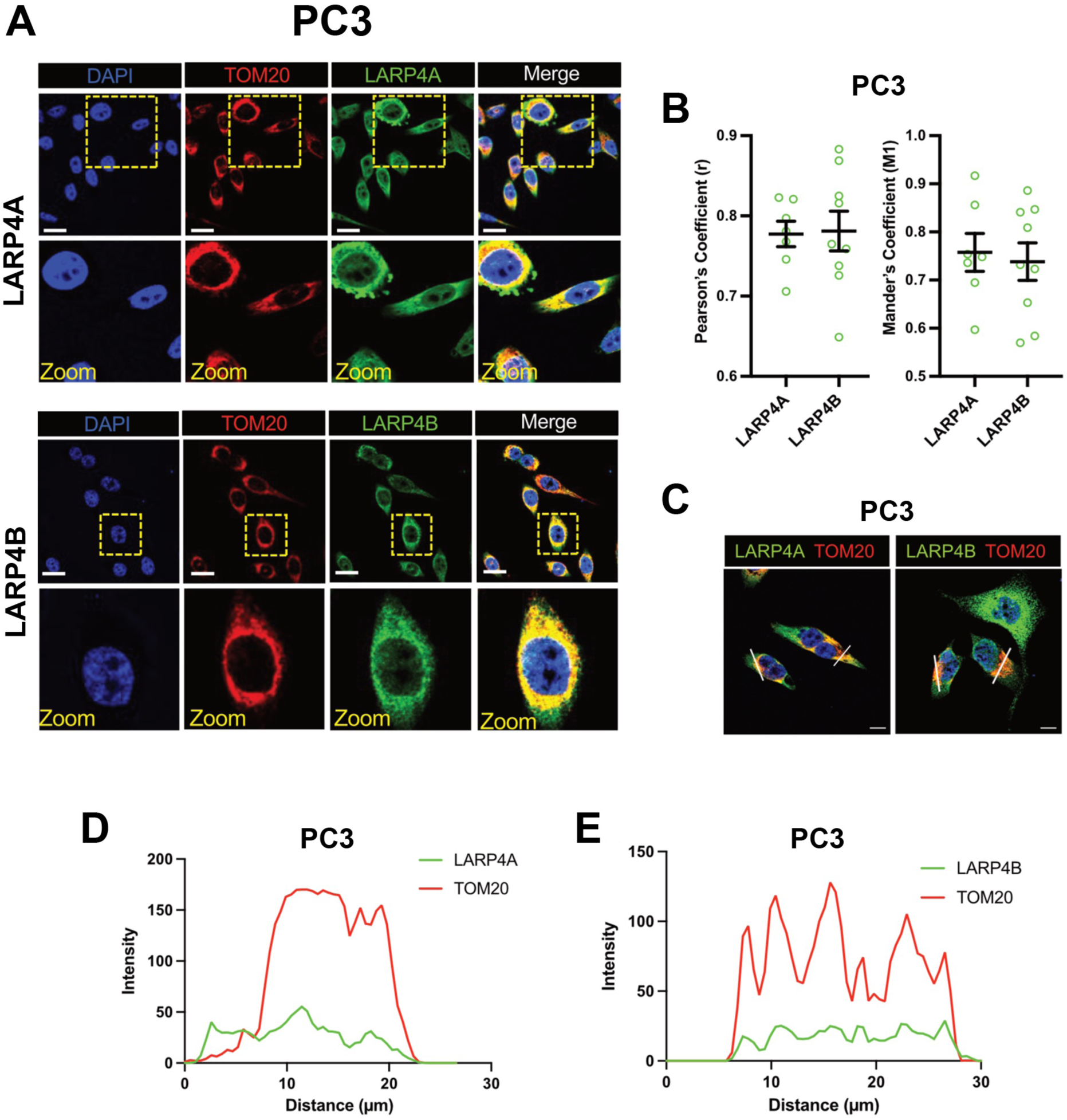
Expression and localisation of LARP4A and LARP4B in PC3 cell mitochondria. (A) Immunofluorescence detection of TOM20 and co-localisation with LARP4A (*top*) and LARP4B (*bottom*) with DAPI counterstain in PC3 cells. Scale bars, 25µm. Areas in yellow squares are depicted at higher magnification (zoom) in lower panels. (B) Pearson’s and Mander’s coefficient analyses showing a similar degree of LARP4B/TOM20 co-localisation compared with LARP4A/TOM20 in PC3 cells. The data represent the mean ± SEM, n=6 from three independent experiments. p-values are indicated. (C-E) Fluorescent intensity line plot profiling demonstrating co-localisation of LARP4A and LARP4B with the TOM20 mitochondrial marker in PC3 cells. White lines in (C) depict the areas used for LARP4A/TOM20 (D) and LARP4B/TOM20 (E) line plot analyses. Scale bars in (C), 10µm.

**Figure S7.**
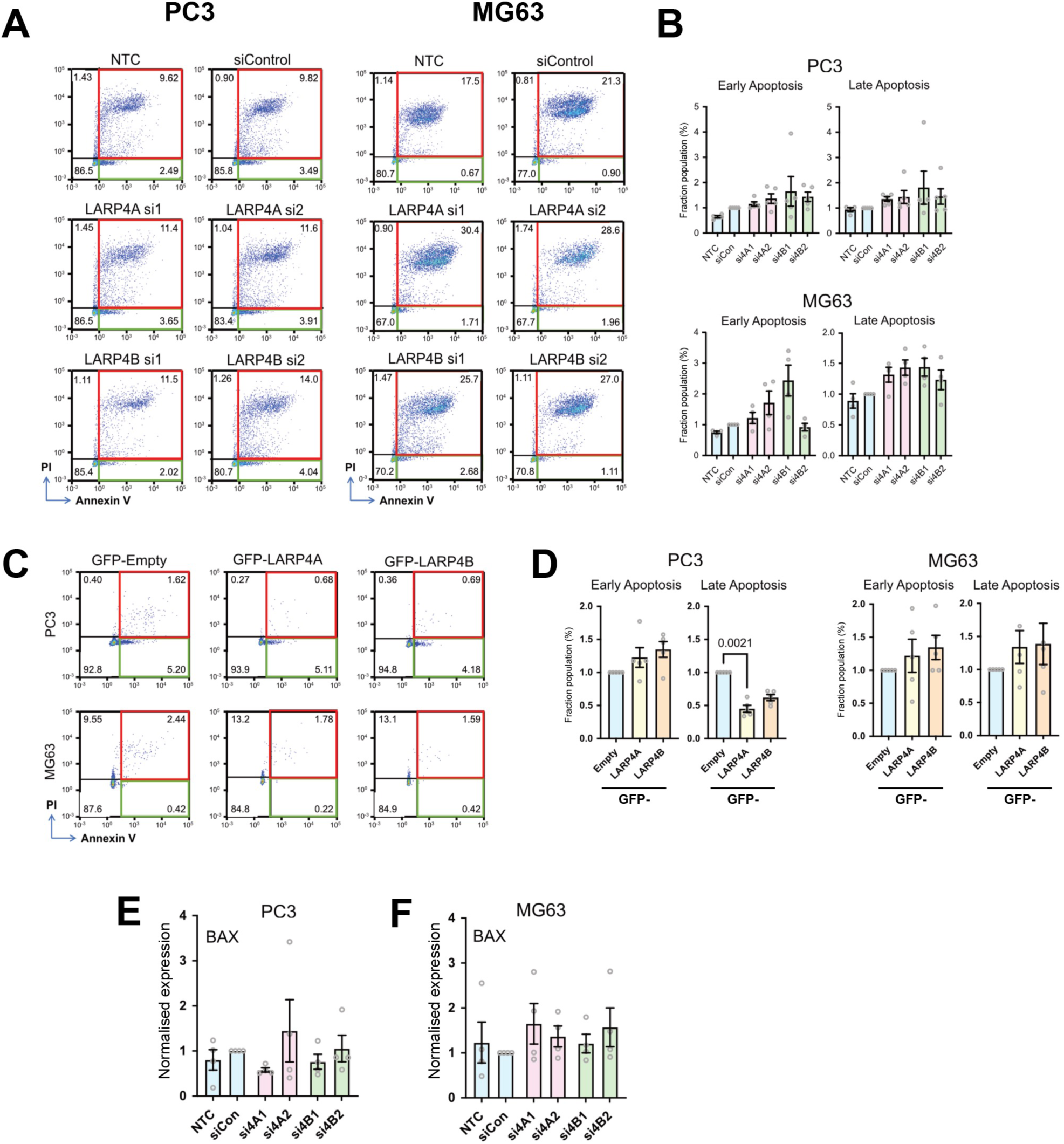
LARP4A or LARP4B silencing does not affect apoptosis of PC3 or MG63 cells. (A) Flow cytometry/Annexin V analysis of PC3 and MG63 cells either non-transfected (NTC), or transfected with non-targeting siControl and siRNAs targeting LARP4A (LARP4Asi1, LARP4Asi2) and LARP4B (LARP4Bsi1, LARP4Bsi2) as indicated. Red boxes denote cells in late apoptosis (Annexin V+/PI+), green boxes denote cells in early apoptosis (Annexin V+/PI-). The percentage population in each cell stage are shown within each quadrant. (B) Quantification of the percentage population of PC3 (*top*) or MG63 (*bottom*) control (NTC, siCon), siLARP4A (siA1, siA2) and siLARP4B (siB1, siB2) knockdown cells in early and late stages of apoptosis. The data represent the mean ± SEM, n=4 independent experiments. (C) Flow cytometry/Annexin V analysis of PC3 and MG63 cells overexpressing LARP4A or LARP4B following transfection with empty GFP (Empty), GFP-LARP4A or GFP-LARP4B plasmids. Red boxes denote cells in late apoptosis (Annexin V+/PI+), green boxes denote cells in early apoptosis (Annexin V+/PI-). The percentage population in each cell stage are shown within each quadrant. (D) Quantification of the percentage population of PC3 (*left*) or MG63 (*right*) control (Empty), GFP-LARP4A and GFP-LARP4B overexpressing cells in early and late stages of apoptosis. The data represent the mean ± SEM, n=3 independent experiments. (E and F) Expression of *BAX* mRNA expression in PC3 (E) and MG63 (F) control (NTC, siCon), siLARP4A (si4A1, si4A2) and siLARP4B (si4B1, si4B2) knockdown cells. Relative mRNA expression is normalised to ý-actin as a housekeeping gene and to siControl cells. The data represent the mean ± SEM, n=3.

**Figure S8.**
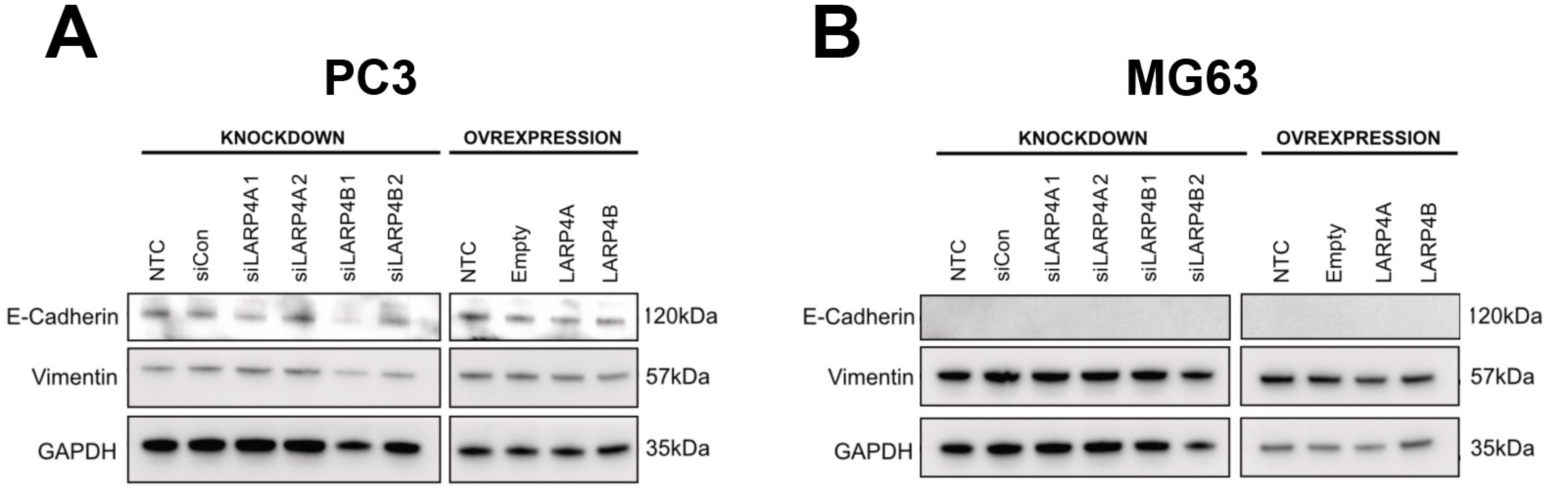
LARP4A and LARP4B do not regulate EMT in PC3 and MG63 cells, and regulate osteosarcoma cell migration. (A and B) Western blot analysis of EMT markers E-Cadherin and Vimentin in PC3 (A) and MG63 (B) cells following either LARP4A or LARP4B knockdown (*left*) or overexpression (*right*). Knockdown cells were transfected with either control siRNAs (NTC, siCon) or siRNAs for LARP4A (siLARP4A1, siLARP4A2) and LARP4B (siLARP4B1, siLARP4B2) as indicated. Overexpression cells were transfected with either Empty or GFP-LARP4A and GRP-LARP4B plasmids as indicated. GAPDH used as a loading control.

**Figure S9.**
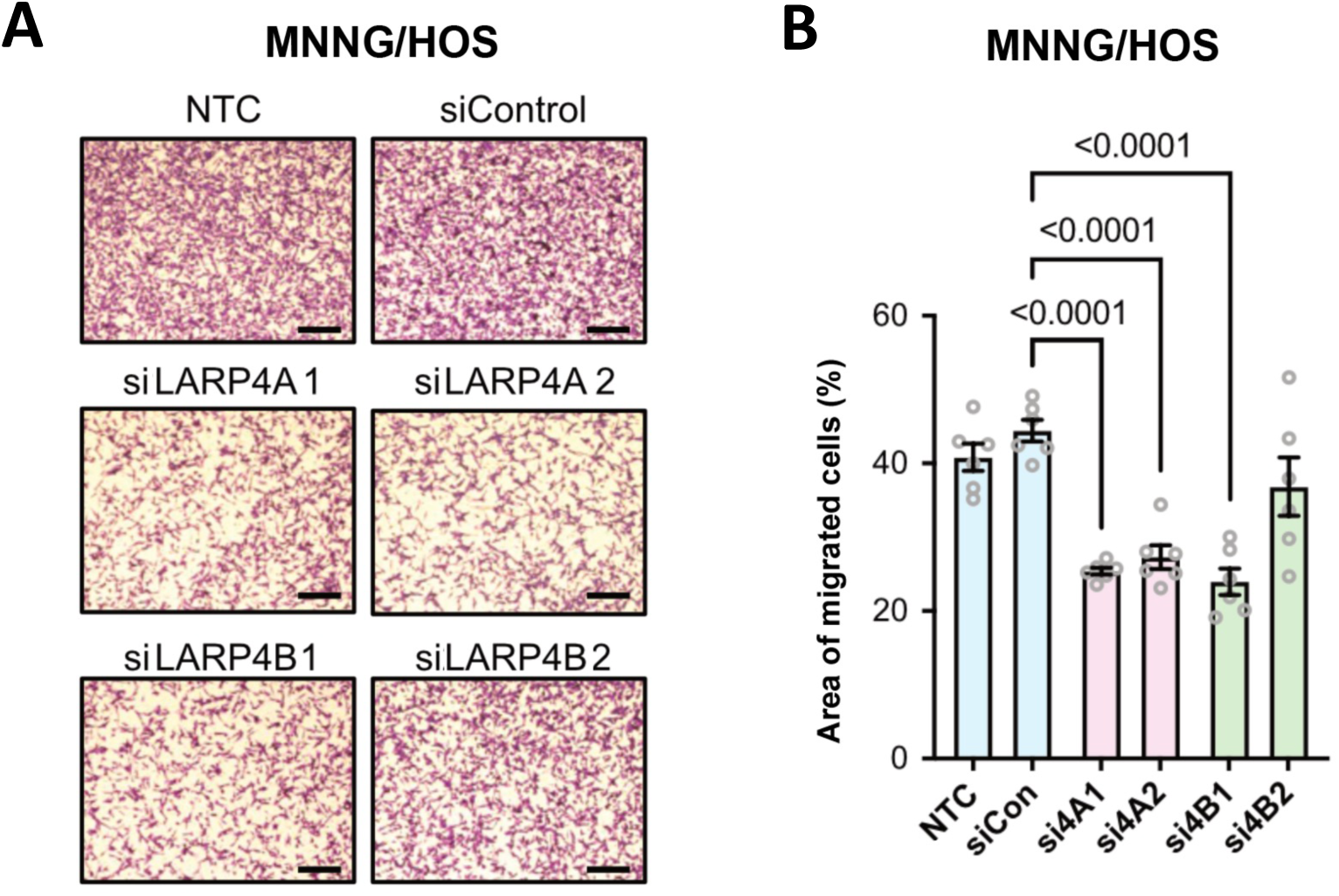
LARP4A and LARP4B regulate cell migration and invasion in MNNG-HOS cells. (A) Brightfield images of transwells stained with crystal violet following a 24h migration assay of MNNG/HOS control (siControl), siLARP4A (4A1, 4A2) or siLARP4B (4B1, 4B2) knockdown cells. Scale bars, 200µm. (B) Quantification of migrated MNNG/HOS cells over 24h of either non-transfected cells (NTC, siCon) or cells transfected with siRNAs targeting LARP4A (siA1, siA2) or LARP4B (siB1, siB2). The data represent the mean ± SEM, n=6 from 3 independent experiments. p-values are indicated.

## REFERENCES

1. Bailey, M.H., Tokheim, C., Porta-Pardo, E., Sengupta, S., Bertrand, D., Weerasinghe, A., Colaprico, A., Wendl, M.C., Kim, J., Reardon, B., et al. (2018). Comprehensive Characterization of Cancer Driver Genes and Mutations. Cell 173, 371–385 e318. 10.1016/j.cell.2018.02.060.

2. Hanahan, D., and Weinberg, R.A. (2011). Hallmarks of cancer: the next generation. Cell 144, 646–674. 10.1016/j.cell.2011.02.013.

3. Schwanhäusser, B., Busse, D., Li, N., Dittmar, G., Schuchhardt, J., Wolf, J., Chen, W., and Selbach, M. (2011). Global quantification of mammalian gene expression control. Nature 473, 337–342. 10.1038/nature10098.

4. Sharp, P.A. (2009). The Centrality of RNA. Cell 136, 577–580. 10.1016/j.cell.2009.02.007.

5. Müller-Mcnicoll, M., and Neugebauer, K.M. (2013). How cells get the message: dynamic assembly and function of mRNA–protein complexes. Nature Reviews Genetics 14, 275–287. 10.1038/nrg3434.

6. Castello, A., Fischer, B., Eichelbaum, K., Horos, R., Beckmann, B.M., Strein, C., Davey, N.E., Humphreys, D.T., Preiss, T., Steinmetz, L.M., et al. (2012). Insights into RNA biology from an atlas of mammalian mRNA-binding proteins. Cell 149, 1393–1406. 10.1016/j.cell.2012.04.031.

7. Van Nostrand, E.L., Freese, P., Pratt, G.A., Wang, X., Wei, X., Xiao, R., Blue, S.M., Chen, J.-Y., Cody, N.A.L., Dominguez, D., et al. (2020). A large-scale binding and functional map of human RNA-binding proteins. Nature 583, 711–719. 10.1038/s41586-020-2077-3.

8. Choi, P.S., and Thomas-Tikhonenko, A. (2021). RNA-binding proteins of COSMIC importance in cancer. J Clin Invest 131. 10.1172/JCI151627.

9. Lukong, K.E., Chang, K.W., Khandjian, E.W., and Richard, S. (2008). RNA-binding proteins in human genetic disease. Trends Genet 24, 416–425. 10.1016/j.tig.2008.05.004.

10. Kang, D., Lee, Y., and Lee, J.-S. (2020). RNA-Binding Proteins in Cancer: Functional and Therapeutic Perspectives. Cancers 12, 2699. 10.3390/cancers12092699.

11. Maraia, R.J., Mattijssen, S., Cruz-Gallardo, I., and Conte, M.R. (2017). The La and related RNA-binding proteins (LARPs): structures, functions, and evolving perspectives. WIREs RNA 8, e1430. 10.1002/wrna.1430.

12. Bousquet-Antonelli, C., and Deragon, J.M. (2009). A comprehensive analysis of the La-motif protein superfamily. RNA 15, 750–764. 10.1261/rna.1478709.

13. Dock-Bregeon, A.C., Lewis, K.A., and Conte, M.R. (2021). The La-related proteins: structures and interactions of a versatile superfamily of RNA-binding proteins. RNA Biol 18, 178–193. 10.1080/15476286.2019.1695712.

14. Lizarrondo, J., Dock-Bregeon, A.-C., Martino, L., and Conte, M.R. (2021). Structural dynamics in the La-module of La-related proteins. RNA Biology 18, 194–206. 10.1080/15476286.2020.1733799.

15. Martino, L., Pennell, S., Kelly, G., Busi, B., Brown, P., Atkinson, R.A., Salisbury, N.J., Ooi, Z.H., See, K.W., Smerdon, S.J., et al. (2015). Synergic interplay of the La motif, RRM1 and the interdomain linker of LARP6 in the recognition of collagen mRNA expands the RNA binding repertoire of the La module. Nucleic Acids Res 43, 645–660. 10.1093/nar/gku1287.

16. Kotik-Kogan, O., Valentine, E.R., Sanfelice, D., Conte, M.R., and Curry, S. (2008). Structural analysis reveals conformational plasticity in the recognition of RNA 3’ ends by the human La protein. Structure 16, 852–862. 10.1016/j.str.2008.02.021.

17. Alfano, C., Sanfelice, D., Babon, J., Kelly, G., Jacks, A., Curry, S., and Conte, M.R. (2004). Structural analysis of cooperative RNA binding by the La motif and central RRM domain of human La protein. Nature structural & molecular biology 11, 323–329.

18. Uchikawa, E., Natchiar, K.S., Han, X., Proux, F., Roblin, P., Zhang, E., Durand, A., Klaholz, B.P., and Dock-Bregeon, A.C. (2015). Structural insight into the mechanism of stabilization of the 7SK small nuclear RNA by LARP7. Nucleic Acids Res 43, 3373–3388. 10.1093/nar/gkv173.

19. Kozlov, G., Mattijssen, S., Jiang, J., Nyandwi, S., Sprules, T., Iben, J.R., Coon, S.L., Gaidamakov, S., Noronha, A.M., Wilds, C.J., et al. (2022). Structural basis of 3’-end poly(A) RNA recognition by LARP1. Nucleic Acids Res 50, 9534–9547. 10.1093/nar/gkac696.

20. Cruz-Gallardo, I., Martino, L., Kelly, G., Atkinson, R.A., Trotta, R., De Tito, S., Coleman, P., Ahdash, Z., Gu, Y., Bui, T.T.T., and Conte, M.R. (2019). LARP4A recognizes polyA RNA via a novel binding mechanism mediated by disordered regions and involving the PAM2w motif, revealing interplay between PABP, LARP4A and mRNA. Nucleic Acids Res 47, 4272–4291. 10.1093/nar/gkz144.

21. Stavraka, C., and Blagden, S. (2015). The La-Related Proteins, a Family with Connections to Cancer. Biomolecules 5, 2701–2722. 10.3390/biom5042701.

22. Grimm, C., Pelz, J.P., Schneider, C., Schaffler, K., and Fischer, U. (2020). Crystal Structure of a Variant PAM2 Motif of LARP4B Bound to the MLLE Domain of PABPC1. Biomolecules 10. 10.3390/biom10060872.

23. Schäffler, K., Schulz, K., Hirmer, A., Wiesner, J., Grimm, M., Sickmann, A., and Fischer, U. (2010). A stimulatory role for the La-related protein 4B in translation. RNA 16, 1488–1499. 10.1261/rna.2146910.

24. Yang, R., Gaidamakov, S.A., Xie, J., Lee, J., Martino, L., Kozlov, G., Crawford, A.K., Russo, A.N., Conte, M.R., Gehring, K., and Maraia, R.J. (2011). La-Related Protein 4 Binds Poly(A), Interacts with the Poly(A)-Binding Protein MLLE Domain via a Variant PAM2w Motif, and Can Promote mRNA Stability. Molecular and Cellular Biology 31, 542–556. 10.1128/mcb.01162-10.

25. Küspert, M., Murakawa, Y., Schaffler, K., Vanselow, J.T., Wolf, E., Juranek, S., Schlosser, A., Landthaler, M., and Fischer, U. (2015). LARP4B is an AU-rich sequence associated factor that promotes mRNA accumulation and translation. RNA 21, 1294–1305. 10.1261/rna.051441.115.

26. Mattijssen, S., Arimbasseri, A.G., Iben, J.R., Gaidamakov, S., Lee, J., Hafner, M., and Maraia, R.J. (2017). LARP4 mRNA codon-tRNA match contributes to LARP4 activity for ribosomal protein mRNA poly(A) tail length protection. eLife 6. 10.7554/elife.28889.

27. Mattijssen, S., Iben, J.R., Li, T., Coon, S.L., and Maraia, R.J. (2020). Single molecule poly(A) tail-seq shows LARP4 opposes deadenylation throughout mRNA lifespan with most impact on short tails. eLife 9. 10.7554/elife.59186.

28. Tian, Y., Zeng, Z., Li, X., Wang, Y., Chen, R., Mattijssen, S., Gaidamakov, S., Wu, Y., Maraia, R.J., Peng, W., and Zhu, J. (2020). Transcriptome-wide stability analysis uncovers LARP4-mediated NFkappaB1 mRNA stabilization during T cell activation. Nucleic Acids Res 48, 8724–8739. 10.1093/nar/gkaa643.

29. Kim, S., Kim, S., Chang, H.R., Kim, D., Park, J., Son, N., Park, J., Yoon, M., Chae, G., Kim, Y.K., et al. (2021). The regulatory impact of RNA-binding proteins on microRNA targeting. Nat Commun 12, 5057. 10.1038/s41467-021-25078-5.

30. Gabrovsek, L., Collins, K.B., Aggarwal, S., Saunders, L.M., Lau, H.-T., Suh, D., Sancak, Y., Trapnell, C., Ong, S.-E., Smith, F.D., and Scott, J.D. (2020). A-kinase-anchoring protein 1 (dAKAP1)-based signaling complexes coordinate local protein synthesis at the mitochondrial surface. Journal of Biological Chemistry 295, 10749–10765. 10.1074/jbc.ra120.013454.

31. Funakoshi, M., Tsuda, M., Muramatsu, K., Hatsuda, H., Morishita, S., and Aigaki, T. (2018). Overexpression of Larp4B downregulates dMyc and reduces cell and organ sizes in Drosophila. Biochem Biophys Res Commun 497, 762–768. 10.1016/j.bbrc.2018.02.148.

32. Deragon, J.M. (2021). Distribution, organization an evolutionary history of La and LARPs in eukaryotes. RNA Biol 18, 159–167. 10.1080/15476286.2020.1739930.

33. Merret, R., Martino, L., Bousquet-Antonelli, C., Fneich, S., Descombin, J., Billey, É., Conte, M.R., and Deragon, J.-M. (2013). The association of a La module with the PABP-interacting motif PAM2 is a recurrent evolutionary process that led to the neofunctionalization of La-related proteins. RNA 19, 36–50. 10.1261/rna.035469.112.

34. Bai, S.W., Herrera-Abreu, M.T., Rohn, J.L., Racine, V., Tajadura, V., Suryavanshi, N., Bechtel, S., Wiemann, S., Baum, B., and Ridley, A.J. (2011). Identification and characterization of a set of conserved and new regulators of cytoskeletal organization, cell morphology and migration. BMC biology 9, 1–18.

35. Egiz, M., Usui, T., Ishibashi, M., Zhang, X., Shigeta, S., Toyoshima, M., Kitatani, K., and Yaegashi, N. (2019). La-Related Protein 4 as a Suppressor for Motility of Ovarian Cancer Cells. Tohoku J Exp Med 247, 59–67. 10.1620/tjem.247.59.

36. Seetharaman, S., Flemyng, E., Shen, J., Conte, M.R., and Ridley, A.J. (2016). The RNA-binding protein LARP4 regulates cancer cell migration and invasion. Cytoskeleton 73, 680–690. 10.1002/cm.21336.

37. Koso, H., Takeda, H., Yew, C.C., Ward, J.M., Nariai, N., Ueno, K., Nagasaki, M., Watanabe, S., Rust, A.G., Adams, D.J., et al. (2012). Transposon mutagenesis identifies genes that transform neural stem cells into glioma-initiating cells. Proc Natl Acad Sci U S A 109, E2998–3007. 10.1073/pnas.1215899109.

38. Koso, H., Yi, H., Sheridan, P., Miyano, S., Ino, Y., Todo, T., and Watanabe, S. (2016). Identification of RNA-Binding Protein LARP4B as a Tumor Suppressor in Glioma. Cancer Res 76, 2254–2264. 10.1158/0008-5472.CAN-15-2308.

39. Zhang, Y., Peng, L., Hu, T., Wan, Y., Ren, Y., Zhang, J., Wang, X., Zhou, Y., Yuan, W., Wang, Q., et al. (2015). La-related protein 4B maintains murine MLL-AF9 leukemia stem cell self-renewal by regulating cell cycle progression. Experimental Hematology 43, 309–318.e302. 10.1016/j.exphem.2014.12.003.

40. Li, Y., Jiao, Y., Li, Y., and Liu, Y. (2019). Expression of La Ribonucleoprotein Domain Family Member 4B (LARP4B) in Liver Cancer and Their Clinical and Prognostic Significance. Disease Markers 2019, 1–13. 10.1155/2019/1569049.

41. Takeda, H., Wei, Z., Koso, H., Rust, A.G., Yew, C.C.K., Mann, M.B., Ward, J.M., Adams, D.J., Copeland, N.G., and Jenkins, N.A. (2015). Transposon mutagenesis identifies genes and evolutionary forces driving gastrointestinal tract tumor progression. Nature Genetics 47, 142–150. 10.1038/ng.3175.

42. Bard-Chapeau, E.A., Nguyen, A.T., Rust, A.G., Sayadi, A., Lee, P., Chua, B.Q., New, L.S., de Jong, J., Ward, J.M., Chin, C.K., et al. (2014). Transposon mutagenesis identifies genes driving hepatocellular carcinoma in a chronic hepatitis B mouse model. Nat Genet 46, 24–32. 10.1038/ng.2847.

43. Rahrmann, E.P., Watson, A.L., Keng, V.W., Choi, K., Moriarity, B.S., Beckmann, D.A., Wolf, N.K., Sarver, A., Collins, M.H., Moertel, C.L., et al. (2013). Forward genetic screen for malignant peripheral nerve sheath tumor formation identifies new genes and pathways driving tumorigenesis. Nature Genetics 45, 756–766. 10.1038/ng.2641.

44. Wu, X., Northcott, P.A., Dubuc, A., Dupuy, A.J., Shih, D.J.H., Witt, H., Croul, S., Bouffet, E., Fults, D.W., Eberhart, C.G., et al. (2012). Clonal selection drives genetic divergence of metastatic medulloblastoma. Nature 482, 529–533. 10.1038/nature10825.

45. Hu, Z., Yau, C., and Ahmed, A.A. (2017). A pan-cancer genome-wide analysis reveals tumour dependencies by induction of nonsense-mediated decay. Nat Commun 8, 15943. 10.1038/ncomms15943.

46. Grigoriadis, A.E., Schellander, K., Wang, Z.Q., and Wagner, E.F. (1993). Osteoblasts are target cells for transformation in c-fos transgenic mice. J Cell Biol 122, 685–701. 10.1083/jcb.122.3.685.

47. Weekes, D., Kashima, T.G., Zandueta, C., Perurena, N., Thomas, D.P., Sunters, A., Vuillier, C., Bozec, A., El-Emir, E., Miletich, I., et al. (2016). Regulation of osteosarcoma cell lung metastasis by the c-Fos/AP-1 target FGFR1. Oncogene 35, 2852–2861. 10.1038/onc.2015.344.

48. Zeng, W.Z.D., Glicksberg, B.S., Li, Y., and Chen, B. (2019). Selecting precise reference normal tissue samples for cancer research using a deep learning approach. BMC Medical Genomics 12. 10.1186/s12920-018-0463-6.

49. Tattersall, L., Shah, K.M., Lath, D.L., Singh, A., Down, J.M., De Marchi, E., Williamson, A., Di Virgilio, F., Heymann, D., Adinolfi, E., et al. (2021). The P2RX7B splice variant modulates osteosarcoma cell behaviour and metastatic properties. J Bone Oncol 31, 100398. 10.1016/j.jbo.2021.100398.

50. Massey, A.J. (2015). Multiparametric Cell Cycle Analysis Using the Operetta High-Content Imager and Harmony Software with PhenoLOGIC. PLoS One 10, e0134306. 10.1371/journal.pone.0134306.

51. Yang, C.-C., Kato, H., Shindo, M., and Masai, H. (2019). Cdc7 activates replication checkpoint by phosphorylating the Chk1-binding domain of Claspin in human cells. eLife 8. 10.7554/elife.50796.

52. Denechaud, P.D., Fajas, L., and Giralt, A. (2017). E2F1, a Novel Regulator of Metabolism. Front Endocrinol (Lausanne) 8, 311. 10.3389/fendo.2017.00311.

53. Uxa, S., Castillo-Binder, P., Kohler, R., Stangner, K., Müller, G.A., and Engeland, K. (2021). Ki-67 gene expression. Cell Death & Differentiation 28, 3357–3370. 10.1038/s41418-021-00823-x.

54. Misra, S., Sharma, S., Agarwal, A., Khedkar, S.V., Tripathi, M.K., Mittal, M.K., and Chaudhuri, G. (2010). Cell cycle-dependent regulation of the bi-directional overlapping promoter of human BRCA2/ZAR2 genes in breast cancer cells. Molecular Cancer 9, 50. 10.1186/1476-4598-9-50.

55. Lewis, B.M., Cho, C.Y., Her, H.-L., Hunter, T., and Yeo, G.W. (2022). LARP4 Is an RNA-Binding Protein That Binds Nuclear-Encoded Mitochondrial mRNAs To Promote Mitochondrial Function. Cold Spring Harbor Laboratory.

56. Ben-Salem, S., Venkadakrishnan, V.B., and Heemers, H.V. (2021). Novel insights in cell cycle dysregulation during prostate cancer progression. Endocr Relat Cancer 28, R141–R155. 10.1530/ERC-20-0517.

57. Cheng, L., Wang, C., and Jing, J. (2016). Cell Cycle Kinases in Osteosarcoma: Potential for Therapeutic Intervention. Curr Pharm Des 22, 4830–4834. 10.2174/1381612822666160512151028.

58. Bavetsias, V., and Linardopoulos, S. (2015). Aurora Kinase Inhibitors: Current Status and Outlook. Front Oncol 5, 278. 10.3389/fonc.2015.00278.

59. Zhu, L.B., Jiang, J., Zhu, X.P., Wang, T.F., Chen, X.Y., Luo, Q.F., Shu, Y., Liu, Z.L., and Huang, S.H. (2014). Knockdown of Aurora-B inhibits osteosarcoma cell invasion and migration via modulating PI3K/Akt/NF-kappaB signaling pathway. Int J Clin Exp Pathol 7, 3984–3991.

60. Chieffi, P., Cozzolino, L., Kisslinger, A., Libertini, S., Staibano, S., Mansueto, G., De Rosa, G., Villacci, A., Vitale, M., Linardopoulos, S., et al. (2006). Aurora B expression directly correlates with prostate cancer malignancy and influence prostate cell proliferation. Prostate 66, 326–333. 10.1002/pros.20345.

61. Kansara, M., Teng, M.W., Smyth, M.J., and Thomas, D.M. (2014). Translational biology of osteosarcoma. Nat Rev Cancer 14, 722–735. 10.1038/nrc3838.

62. Nyquist, M.D., Corella, A., Coleman, I., De Sarkar, N., Kaipainen, A., Ha, G., Gulati, R., Ang, L., Chatterjee, P., Lucas, J., et al. (2020). Combined TP53 and RB1 Loss Promotes Prostate Cancer Resistance to a Spectrum of Therapeutics and Confers Vulnerability to Replication Stress. Cell Reports 31, 107669. 10.1016/j.celrep.2020.107669.

63. Zhang, W., and Zhang, K. (2022). A transcriptomic signature for prostate cancer relapse prediction identified from the differentially expressed genes between TP53 mutant and wild-type tumors. Scientific Reports 12. 10.1038/s41598-022-14436-y.

64. Bisogno, L.S., and Keene, J.D. (2018). RNA regulons in cancer and inflammation. Curr Opin Genet Dev 48, 97–103. 10.1016/j.gde.2017.11.004.

65. Blackinton, J.G., and Keene, J.D. (2014). Post-transcriptional RNA regulons affecting cell cycle and proliferation. Semin Cell Dev Biol 34, 44–54. 10.1016/j.semcdb.2014.05.014.

66. Epstein, T., Xu, L., Gillies, R.J., and Gatenby, R.A. (2014). Separation of metabolic supply and demand: aerobic glycolysis as a normal physiological response to fluctuating energetic demands in the membrane. Cancer Metab 2, 7. 10.1186/2049-3002-2-7.

67. Shiraishi, T., Verdone, J.E., Huang, J., Kahlert, U.D., Hernandez, J.R., Torga, G., Zarif, J.C., Epstein, T., Gatenby, R., McCartney, A., et al. (2015). Glycolysis is the primary bioenergetic pathway for cell motility and cytoskeletal remodeling in human prostate and breast cancer cells. Oncotarget 6, 130–143. 10.18632/oncotarget.2766.

68. Zanotelli, M.R., Zhang, J., and Reinhart-King, C.A. (2021). Mechanoresponsive metabolism in cancer cell migration and metastasis. Cell Metabolism 33, 1307–1321. 10.1016/j.cmet.2021.04.002.

69. Gilbertson, S., Federspiel, J.D., Hartenian, E., Cristea, I.M., and Glaunsinger, B. (2018). Changes in mRNA abundance drive shuttling of RNA binding proteins, linking cytoplasmic RNA degradation to transcription. Elife 7. 10.7554/eLife.37663.

70. Conrad, T., Albrecht, A.S., de Melo Costa, V.R., Sauer, S., Meierhofer, D., and Orom, U.A. (2016). Serial interactome capture of the human cell nucleus. Nat Commun 7, 11212. 10.1038/ncomms11212.

71. Denisenko, T.V., Gorbunova, A.S., and Zhivotovsky, B. (2019). Mitochondrial Involvement in Migration, Invasion and Metastasis. Front Cell Dev Biol 7, 355. 10.3389/fcell.2019.00355.

72. Valcarcel-Jimenez, L., Gaude, E., Torrano, V., Frezza, C., and Carracedo, A. (2017). Mitochondrial Metabolism: Yin and Yang for Tumor Progression. Trends in Endocrinology & Metabolism 28, 748–757. 10.1016/j.tem.2017.06.004.

73. Yadav, T., Gau, D., and Roy, P. (2022). Mitochondria-actin cytoskeleton crosstalk in cell migration. J Cell Physiol 237, 2387–2403. 10.1002/jcp.30729.

74. Zhang, J., Miao, X., Wu, T., Jia, J., and Cheng, X. (2021). Development and Validation of Ten-RNA Binding Protein Signature Predicts Overall Survival in Osteosarcoma. Front Mol Biosci 8, 751842. 10.3389/fmolb.2021.751842.

75. Gao, L., Meng, J., Zhang, Y., Gu, J., Han, Z., Wang, X., and Gao, S. (2020). Development and validation of a six-RNA binding proteins prognostic signature and candidate drugs for prostate cancer. Genomics 112, 4980–4992. 10.1016/j.ygeno.2020.08.034.

76. Bankhead, P., Loughrey, M.B., Fernandez, J.A., Dombrowski, Y., McArt, D.G., Dunne, P.D., McQuaid, S., Gray, R.T., Murray, L.J., Coleman, H.G., et al. (2017). QuPath: Open source software for digital pathology image analysis. Sci Rep 7, 16878. 10.1038/s41598-017-17204-5.

77. Bolte, S., and Cordelieres, F.P. (2006). A guided tour into subcellular colocalization analysis in light microscopy. J Microsc 224, 213–232. 10.1111/j.1365-2818.2006.01706.x.

78. Schneider, C.A., Rasband, W.S., and Eliceiri, K.W. (2012). NIH Image to ImageJ: 25 years of image analysis. Nature Methods 9, 671–675. 10.1038/nmeth.2089.

79. Goldman, M., Craft, B., Hastie, M., Repečka, K., McDade, F., Kamath, A., Banerjee, A., Luo, Y., Rogers, D., and Brooks, A.N. (2018). The UCSC Xena platform for public and private cancer genomics data visualization and interpretation. biorxiv, 326470.

80. Goldman, M.J., Craft, B., Hastie, M., Repecka, K., McDade, F., Kamath, A., Banerjee, A., Luo, Y., Rogers, D., Brooks, A.N., et al. (2020). Visualizing and interpreting cancer genomics data via the Xena platform. Nat Biotechnol 38, 675–678. 10.1038/s41587-020-0546-8.

81. Kent, W.J., Sugnet, C.W., Furey, T.S., Roskin, K.M., Pringle, T.H., Zahler, A.M., and Haussler, D. (2002). The human genome browser at UCSC. Genome Res 12, 996–1006. 10.1101/gr.229102.

82. Andrews, S. (n.d.). FastQC: A Quality Control Tool for High Throughput Sequence Data [Online]. http://www.bioinformatics.babraham.ac.uk/projects/fastqc/.

83. Kim, D., Langmead, B., and Salzberg, S.L. (2015). HISAT: a fast spliced aligner with low memory requirements. Nat Methods 12, 357–360. 10.1038/nmeth.3317.

84. Liao, Y., Smyth, G.K., and Shi, W. (2014). featureCounts: an efficient general purpose program for assigning sequence reads to genomic features. Bioinformatics 30, 923–930. 10.1093/bioinformatics/btt656.

85. Liu, R., Holik, A.Z., Su, S., Jansz, N., Chen, K., Leong, H.S., Blewitt, M.E., Asselin-Labat, M.L., Smyth, G.K., and Ritchie, M.E. (2015). Why weight? Modelling sample and observational level variability improves power in RNA-seq analyses. Nucleic Acids Res 43, e97. 10.1093/nar/gkv412.

86. Robinson, M.D., McCarthy, D.J., and Smyth, G.K. (2010). edgeR: a Bioconductor package for differential expression analysis of digital gene expression data. Bioinformatics 26, 139–140. 10.1093/bioinformatics/btp616.

87. Chen, E.Y., Tan, C.M., Kou, Y., Duan, Q., Wang, Z., Meirelles, G.V., Clark, N.R., and Ma’ayan, A. (2013). Enrichr: interactive and collaborative HTML5 gene list enrichment analysis tool. BMC Bioinformatics 14, 128. 10.1186/1471-2105-14-128.

88. Kuleshov, M.V., Jones, M.R., Rouillard, A.D., Fernandez, N.F., Duan, Q., Wang, Z., Koplev, S., Jenkins, S.L., Jagodnik, K.M., Lachmann, A., et al. (2016). Enrichr: a comprehensive gene set enrichment analysis web server 2016 update. Nucleic Acids Res 44, W90–97. 10.1093/nar/gkw377.

89. Xie, Z., Bailey, A., Kuleshov, M.V., Clarke, D.J.B., Evangelista, J.E., Jenkins, S.L., Lachmann, A., Wojciechowicz, M.L., Kropiwnicki, E., Jagodnik, K.M., et al. (2021). Gene Set Knowledge Discovery with Enrichr. Curr Protoc 1, e90. 10.1002/cpz1.90.

